# Bispecific antibody shuttles targeting CD98hc mediate efficient and long-lived brain delivery of IgGs

**DOI:** 10.1101/2023.04.29.538811

**Authors:** Ghasidit Pornnoppadol, Layne G. Bond, Michael J. Lucas, Jennifer M. Zupancic, Yun-Huai Kuo, Boya Zhang, Colin F. Greineder, Peter M. Tessier

## Abstract

The inability of antibodies and other biologics to penetrate the blood-brain barrier (BBB) is a key limitation to their use in diagnostic, imaging, and therapeutic applications. One promising strategy is to deliver IgGs using a bispecific BBB shuttle, which involves fusing an IgG with a second affinity ligand that engages a cerebrovascular endothelial target and facilitates transport across the BBB. Nearly all prior efforts have focused on the transferrin receptor (TfR-1) as the prototypical endothelial target despite inherent delivery and safety challenges. Here we report bispecific antibody shuttles that engage CD98hc (also known as 4F2 and SLC3A2), the heavy chain of the large neutral amino acid transporter (LAT1), and efficiently transport IgGs into the brain parenchyma. Notably, CD98hc shuttles lead to much longer-lived brain retention of IgGs than TfR-1 shuttles while enabling more specific brain targeting due to limited CD98hc engagement in the brain parenchyma. We demonstrate the broad utility of the CD98hc shuttles by reformatting three existing IgGs as CD98hc bispecific shuttles and delivering them to the mouse brain parenchyma that either agonize a neuronal receptor (TrkB) or target other endogenous antigens on specific types of brain cells (neurons and astrocytes).

## INTRODUCTION

The brain is highly vascularized, and its hundreds of miles of blood vessels provide an incredible opportunity to, in principle, deliver any molecule of interest to every brain region. However, in practice, the blood-brain barrier (BBB) strictly regulates molecular transport into the parenchyma, only allowing entry to select nutrients and other biomolecules needed for essential cellular functions^1, 2^. Most biologics, ranging from small peptides to large proteins like IgGs, are largely excluded by the BBB. Even in rare cases where specific IgGs accumulate at levels greater than non-specific uptake (∼0.01-0.1% of injected dose), the enhanced levels are modest and the transport mechanism(s) are typically unclear, making it difficult to extrapolate from animal studies to human trials^3, 4^.

While many strategies for brain delivery of IgGs have been proposed, the leading approach is to use an affinity-based shuttle, which engages a cerebrovascular target on the luminal surface of endothelial cells and facilitates transport across the BBB^5–8^. After nearly two decades of research, the first therapeutics using this strategy only recently entered clinical trials^9^. This slow progress may be partially attributed to the nearly exclusive focus on the transferrin receptor (TfR-1) as the prototypical endothelial target^10–12^ and several challenges specific to this iron-transport protein^13^. First, TfR-1 expression is not exclusively endothelial and is, in fact, used as a marker of immature erythrocytes^13^. Second, binding of antibodies and other affinity ligands to TfR-1 can result in lysosomal degradation of both the receptor and bound antibody, leading to decreased receptor levels and suppression of BBB shuttling^7, 8^. Even more complex, the process appears to depend on the affinity, valency, and epitope of TfR-1 antibodies, and conflicting experimental results have confounded attempts to define these relationships^4, 7, 13, 14^. As a result, there is no consensus as to the optimal construct for maximizing overall brain uptake, parenchymal delivery, and retention of shuttled cargo in the brain.

Given these potential issues with TfR-1 mediated shuttling, there continues to be great interest in identifying other BBB receptors that may enable greater parenchyma delivery or different kinetics of BBB transport and brain retention. One of the most promising targets is CD98hc, the heavy chain of the large neutral amino acid transporter (LAT1)^15, 16^. CD98hc is highly expressed on both mouse and human brain endothelium and is present on both sides of the BBB. A 2016 study demonstrated the potential of CD98hc-mediated antibody shuttling, showing that it was capable of achieving greater brain uptake than that typically seen with TfR-1 shuttles in mice^17^. A more recent (2022) study investigated CD98hc-mediated brain delivery of IgGs in cynomolgus monkeys^18^.

These previous studies^17, 18^, while groundbreaking, have several limitations that are the focus of this work. First, the 2016 report only demonstrates brain shuttling of antibody fragments (Fabs) using a so-called ‘1×1 shuttle’ format – i.e., one Fab arm against a brain target (β-secretase) and the other Fab arm against CD98hc^17^. While useful in terms of its monovalent binding to the cerebrovascular target (e.g., CD98hc)^4^, this format is incapable of delivering existing IgGs as reformatted CD98hc shuttles. This is critical, as reformatting bivalent IgGs into monovalent Fabs for incorporation into a 1×1 shuttle inevitably results in loss of binding affinity and avidity, which is likely to compromise biological function in the brain parenchyma. While the 2022 study addresses this limitation, the reported CD98hc shuttles cannot be used in mice as they are not cross-reactive with mouse CD98hc^18^. Second, the investigators in both studies did not publish the CD98hc antibody sequences, preventing validation of their results or use of the reported shuttles for brain delivery of other antibodies. Third, target engagement within the brain was demonstrated for only two antigens (β-secretase and tau), leaving unanswered questions about the range of parenchymal targets that may be accessed via these approaches. Fourth, it was not evaluated in either study if CD98hc shuttles engaged CD98hc in the brain parenchyma without engagement of a primary brain target. This is notable because the widespread uptake of bispecific antibodies into brain cells, as is commonly observed for TfR-1 shuttles^7, 9, 19, 20^, can limit the ability of bispecific antibodies to selectively bind a second target in the brain parenchyma, such as extracellular targets or specific cell types. Finally, both studies reported little comparison of CD98hc versus TfR-1 shuttles at different doses, such as tracer and saturating doses, limiting understanding of their differences in transport capacity or kinetics.

In this study, we have sought to address each of these limitations. First, we report a 2×1 CD98hc shuttle that efficiently delivers IgGs to the brain while maintaining monovalent engagement of CD98hc via a single-chain antibody (scFv) fused to the C-terminus of one of the two IgG heavy chains. Second, we generated our CD98hc shuttle using an independently validated antibody specific for mouse CD98hc with a publicly available sequence^21^ to enable others to reproduce our findings and use our technology for brain delivery of other IgGs. Third, we demonstrate parenchymal engagement of multiple endogenous targets in wild-type mice, including a neurotrophin receptor (TrkB) and cell-surface proteins on multiple types of brain cells (e.g., neurons and astrocytes). Fourth, we demonstrate limited CD98hc target engagement in the brain parenchyma and compare this directly to the dissimilar behavior of an equivalent TfR-1 shuttle. Finally, we extensively characterize the kinetics of the CD98hc shuttle, comparing them to an equivalent TfR-1 shuttle both at tracer and therapeutic doses, revealing unique advantages of each transport pathway. Ultimately, we demonstrate that the CD98hc shuttle is capable of efficient delivery of IgGs across the BBB, with remarkable retention within the brain and the ability to access a wide variety of parenchymal targets in a simple and predictable manner.

## RESULTS

### *In vitro* characterization of CD98hc brain shuttles

Toward our goal of delivering IgGs to the brain while minimizing IgG modification, we fused a CD98hc single-chain (scFv) antibody to the C-terminus of one of the IgG heavy chains, using a short flexible linker to separate the domains (**Figs. 1A** and **S1**). This “2×1 format” (i.e., one single chain antibody attached to a bivalent IgG) was chosen to preserve the affinity and avidity of the IgG being shuttled into the brain parenchyma while mediating monovalent, lower affinity binding to CD98hc. Production of heterodimerized antibody was maximized using knob-into-hole pairing technology and an optimized transfection ratio of the two heavy chain plasmids^22, 23^. The CD98hc-binding scFv was derived from a monoclonal antibody whose sequence has been defined and published (**Fig. S1**)^21^.

**Figure 1.**
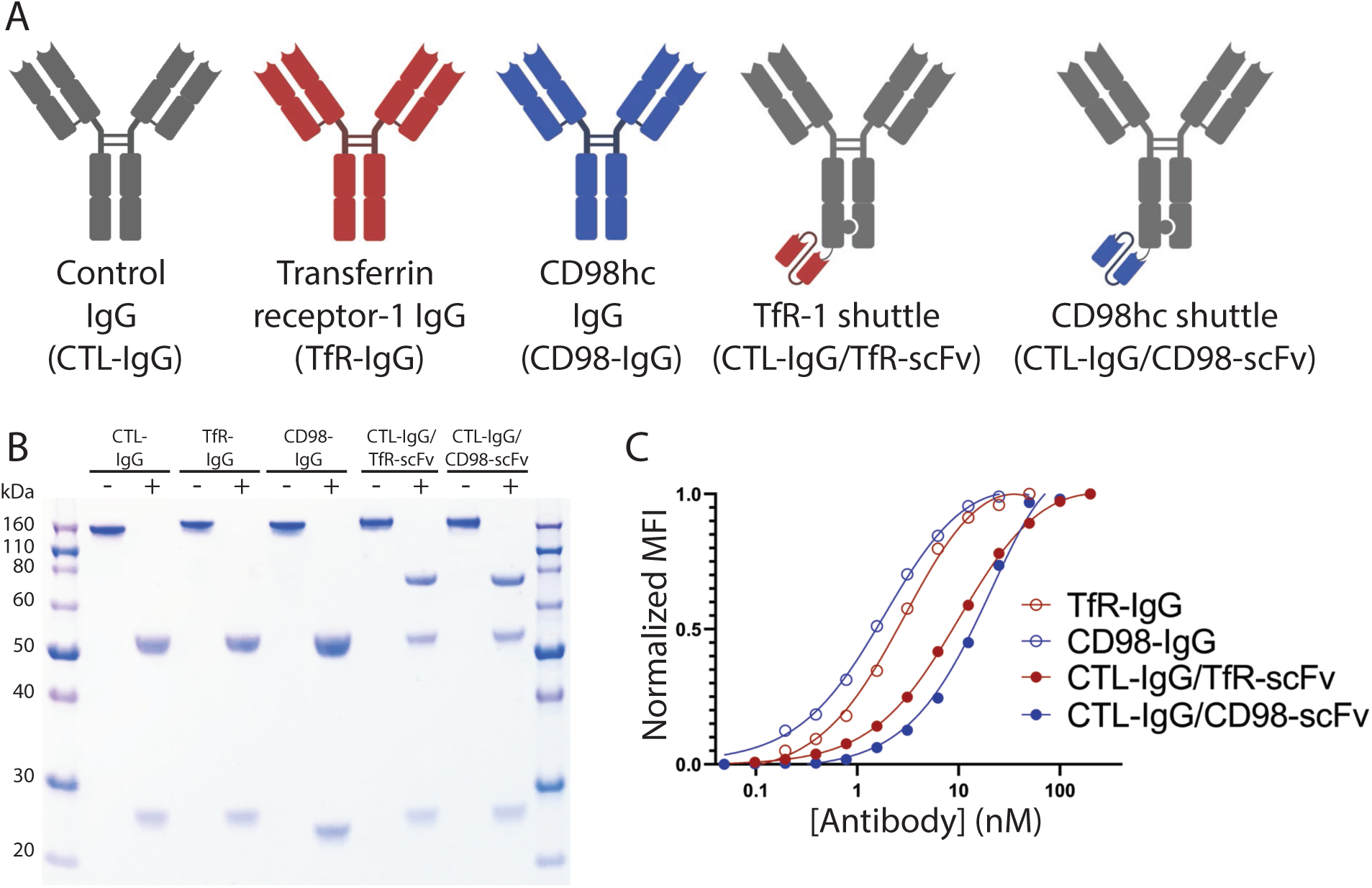
*In vitro* characterization of IgGs and bispecific antibody shuttles. (A) Schematic of the IgGs and bispecific antibodies evaluated in this work. (B) SDS-PAGE analysis of the non-heated/non-reduced (-) and heated/reduced (+) antibodies. Bispecific antibodies display three bands due to the knob-into-hole design, with the heaviest band being the knob portion with the CD98hc single-chain (scFv) antibody. (C) Flow cytometry analysis of IgG and bispecific antibody binding to TfR-1 or CD98hc-expressing cells.

For comparison to the CD98hc shuttle, we generated an analogous 2×1 TfR-1 shuttle using a validated, published, and sequence-defined antibody specific for mouse TfR-1 (clone 8D3) (**Fig. S1**)^24^. Both shuttles were first synthesized with a non-targeted IgG (i.e., specific for human phosphorylated tau without a target in wild-type mice; **Fig. S1)** to enable study of *in vitro* and *in vivo* behavior in the absence of a second target. The shuttles were also engineered with standard Fc mutations to eliminate effector functions^25, 26^. These two shuttles, along with the corresponding non-targeted IgG and two IgGs corresponding to the parental CD98hc and TfR-1 antibodies, were expressed and first purified using Protein A chromatography, which resulted in purification yields of 11-19 mg/L for the IgGs and 5-13 mg/L for the bispecifics. Size-exclusion chromatography (SEC) was also used, as needed, to achieve high purity (>90% monomer, analytical SEC; **Fig. S2**). The proper sizes for the heavy and light chains were also confirmed on a reducing SDS-PAGE gel (**Fig. 1B**). Notably, this analysis revealed the expected three bands for the 2×1 shuttles, with the two heavy chain bands in roughly equal proportion, indicating a high percentage of heterodimer. We used flow cytometry to confirm that the 2×1 shuttles, with only a single binding site for CD98hc or TfR-1, bound to their targets (expressed on human REN cells) with order-of-magnitude lower affinities (EC_50_ of 26.2±3.4 nM for CTL-IgG/CD98-scFv and 10.3±1.4 nM for CTL-IgG/TfR-scFv) relative to their corresponding parental IgGs (EC_50_ of 1.9±0.5 nM for CD98-IgG and 3.1±0.2 nM for TfR-IgG; **Fig. 1C**). Similar binding results were observed using recombinant TfR-1 and CD98hc ectodomains, as the parental IgGs displayed at least order-of-magnitude lower EC_50_s than those for the 2×1 shuttles (**Fig. S3**).

Antibody engagement of TfR-1 is well known to mediate strong receptor internalization^8, 27^. Therefore, we sought to evaluate if our TfR-1 shuttle reduced cell-surface levels of the receptor and if similar behavior is observed for shuttles targeting CD98hc (**Fig. S4**). Using a standard mouse brain endothelial cell line (bEnd.3), we confirmed that both the TfR-1 IgG and shuttle reduced cell-surface levels, which corresponded to 20±3% relative to untreated cells for the IgG and 51±10% for the shuttle. In contrast, the CD98hc IgG and shuttle had more modest impacts on CD98hc cell-surface levels, which corresponded to 75±4% relative to untreated samples for the IgG and 115±2% for the shuttle.

### CD98hc shuttles enable long-lived IgG brain delivery

We next evaluated the pharmacokinetics (PK) of the 2×1 shuttles in wild-type mice. Previous studies of TfR-1-targeted antibodies and shuttles have demonstrated greater brain uptake of high affinity, bivalent binders when given at tracer dose and lower affinity, monovalent binders when given at “saturating” doses^4, 8, 17^. For the latter case, cerebrovascular TfR-1 is completely bound and there is continued brain uptake over a period of hours to days. With this in mind, we performed dose escalation studies to identify the saturating dose of 2×1 shuttles for both CD98hc and TfR-1 (**Fig. S5**). While relatively low doses (0.12 and 0.36 mg/kg) of 2×1 shuttle were not saturating for either target, both targets showed an obvious decrease in brain uptake of radioactive tracer after 1 h at the highest shuttle dose (3.6 mg/kg), indicating at least partial saturation. Interestingly, the pattern of uptake in other organs also indicates saturation of accessible CD98hc in two other major reservoirs – the liver and spleen – at this dose (**Fig. S5**).

Based on these results, we chose 3.6 mg/kg (20 nmol/kg) as the dose for detailed characterization of antibody levels in the brain, blood, and other organs (**Fig. 2**). An equimolar dose of 3 mg/kg (IgGs) and 3.6 mg/kg (shuttles) was used. Consistent with prior reports^4, 8, 28, 29^, the TfR-1 shuttle demonstrated greater brain uptake than its bivalent IgG counterpart, reaching a peak concentration of 5.8±0.1 nM after 8 h (**Fig. 2A**). Despite high initial uptake, the brain concentration of both the TfR-1 shuttle and IgG decreased rapidly with time, falling below 1 nM by 3 days and returning to baseline levels after 5 days. The TfR-1 antibodies showed fast clearance from the blood relative to untargeted IgG (**Fig. 2B**), presumably due to target-mediated drug disposition. Brain:blood ratios were relatively high, peaking after 8 h at 0.16 for the TfR-1 shuttle and 0.5 for the TfR-1 IgG (**Fig. 2C**).

**Figure 2.**
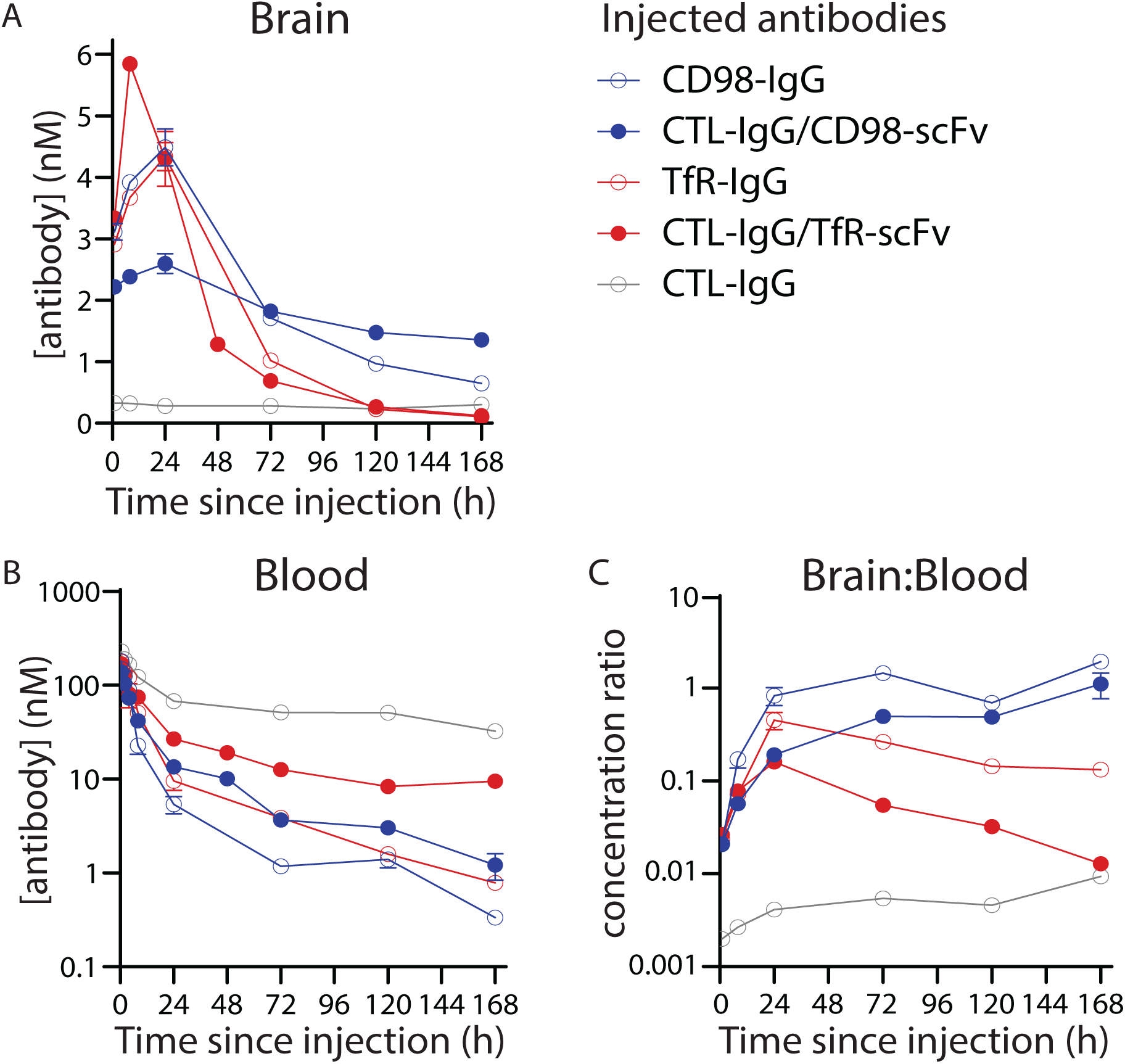
Quantitative pharmacokinetic analysis of CD98hc-and TfR-1-mediated antibody brain uptake. Equimolar doses (20 nmol/kg) of IgGs (3.0 mg/kg) and shuttles (3.6 mg/kg) were radiolabeled with 125I and injected retro-orbitally. Following transcardial perfusion with PBS, concentrations of antibody in the (A) total brain and (B) blood, along with the (C) brain:blood ratios, were determined. n=3-5 mice per antibody group.

Despite the similarity in initial uptake, the CD98hc shuttle displayed distinct brain PK relative to its TfR-1 counterpart (**Fig. 2**). The peak brain concentration was not achieved until 24 h after administration and, in this case, uptake of the bivalent IgG (4.5±0.6 nM) was greater than that of the shuttle (2.6±0.5 nM). However, this finding was dependent on dose, as we observed the opposite behavior at higher doses, namely 9 mg/kg (60 nmol/kg) for CD98hc IgG and 10.8 mg/kg (60 nmol/kg) for CD98hc shuttle (**Fig. S6**). At this higher dose, the CD98hc shuttle reached a higher brain concentration (7.0±0.7 nM) than the CD98hc IgG (5.2±0.4 nM) after 24 h.

Notably, the brain concentration of the CD98hc shuttle remained >1 nM for at least 7 days following a single dose (3.6 mg/kg; **Fig. 2**). As with TfR-1, blood PK appeared to be driven largely by target-mediated effects, with blood concentrations of both CD98hc IgG and shuttle dropping below 10 nM by 24 h post-injection. The combination of blood clearance and sustained brain concentration resulted in impressive brain:blood ratios, which peaked at 1.1 for the CD98hc shuttle and 1.9 for the CD98hc IgG at 7 days post-injection (**Fig. 2C**). Remarkably, the brain:blood ratio for the CD98hc shuttle was ∼110 times higher than that of the TfR-1 shuttle at this time point, highlighting the distinct brain PK of the two shuttles. A statistical analysis of the antibody brain concentrations is provided in **Table S1**, and a summary of the PK in other organs is provided in **Fig. S7**.

Given the potential therapeutic implications of sustained brain delivery for at least one-week post-injection, we sought to confirm the radiotracing results using additional methodologies. First, we injected fluorescently labeled shuttles and IgGs, and evaluated their organ distribution after seven days (**Fig. 3**). Consistent with the radiotracing data, animals injected with a single dose of CD98hc shuttle and IgG had persistent fluorescent signal in their brains for at least 7 days, whereas no signal was seen in those dosed with TfR-1 shuttle, TfR-1 IgG or control IgG. Of note, the fluorescent signal seen in the lungs of mice injected with control IgG likely reflects residual blood content, given its relatively high blood concentration at 7 days (**Fig. 2B**) and variable flushing of the pulmonary circulation when animals are transcardially perfused via the left ventricle. Finally, we performed IgG ELISAs on mice injected with either shuttle and observed brain levels that closely matched the radiotracing results, including results at 3 days that were not statistically different and results at 5 days that displayed minor but statistically significant differences (**Fig. S8**).

**Figure 3.**
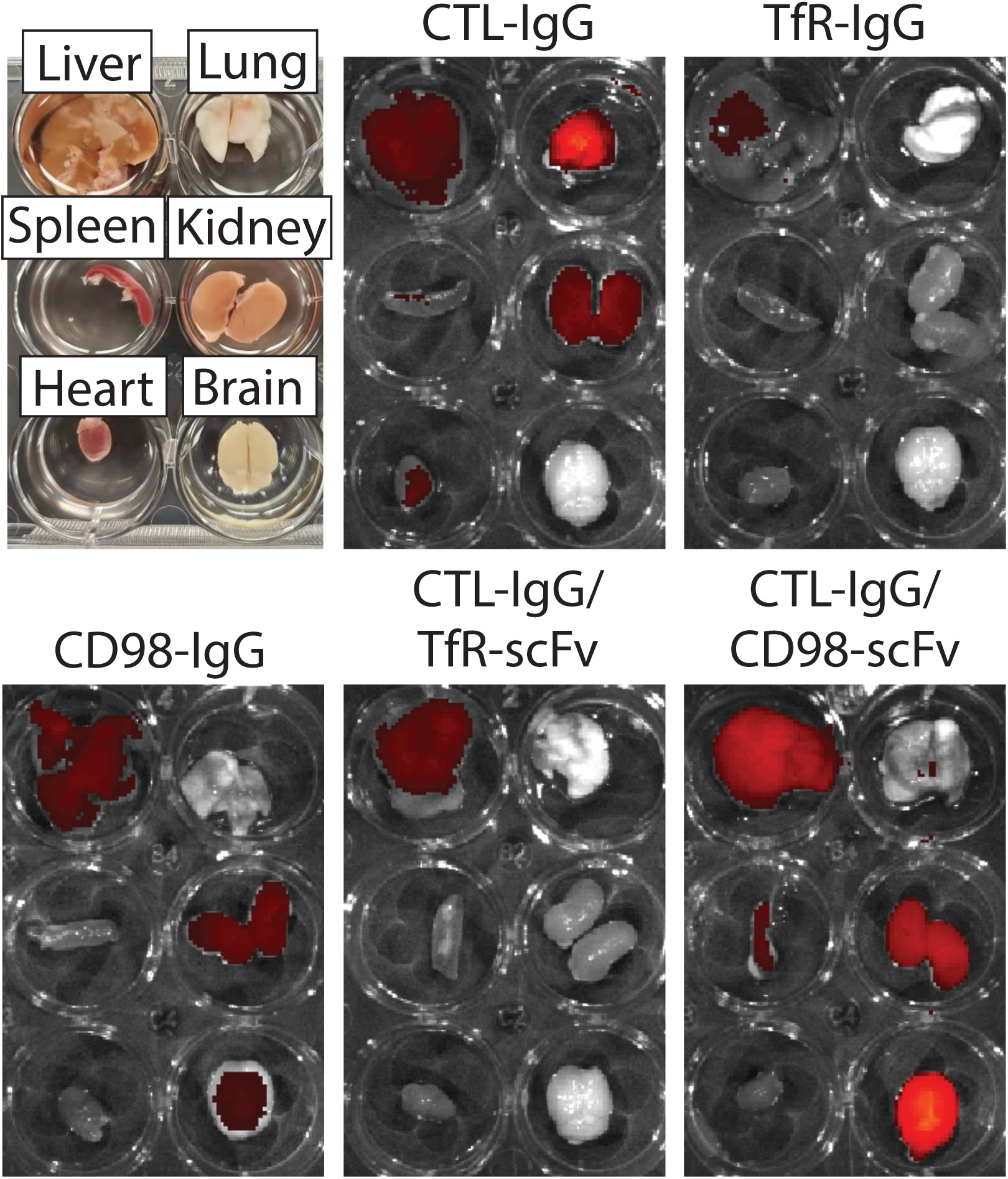
Fluorescent imaging of antibody organ distribution. Equimolar doses (20 nmol/kg) of IgGs (3.0 mg/kg) and shuttles (3.6 mg/kg) were labeled with Alexa Fluor-647 and injected intravenously into wild-type mice. After 7 d, various organs were harvested and imaged with IVIS spectrum. These experiments were repeated in triplicate (n=3), and representative images are shown.

To investigate the distribution of CD98hc-targeted antibodies within brain tissue, we first performed classical fractionation of brain homogenate in animals treated with radiolabeled CD98hc shuttle and IgG. One hour after injection the radioactive shuttles for both cerebrovascular targets were primarily found in the parenchymal fraction, whereas the IgGs were relatively evenly split between capillary and parenchymal fractions (**Fig. S9**). Moreover, we performed fluorescence microscopy on sectioned brain tissue from animals injected with fluorescently labeled shuttles after one to five days post injection (**Fig. 4**). After one day, both shuttles largely localized to blood vessels, as confirmed by co-localization with ZO-1 staining (**Fig. S10**). After three days, fluorescent signal from the TfR-1 shuttle was only weakly associated with blood vessels and appeared within various cell types, including both NeuN-positive neurons and other types of NeuN-negative cells (**Fig. 4**, top). In contrast, the CD98hc shuttle remained largely associated with vessels (**Fig. 4**, bottom). After five days, fluorescent signal from the TfR-1 shuttle was no longer detectable (**Fig. 4**, top), consistent with radiotracing (**Fig. 2**), fluorescence imaging (**Fig. 3**), and ELISA (**Fig. S8**) results, whereas the CD98hc shuttle remained largely associated with vessels (**Fig. 4**, bottom).

**Figure 4.**
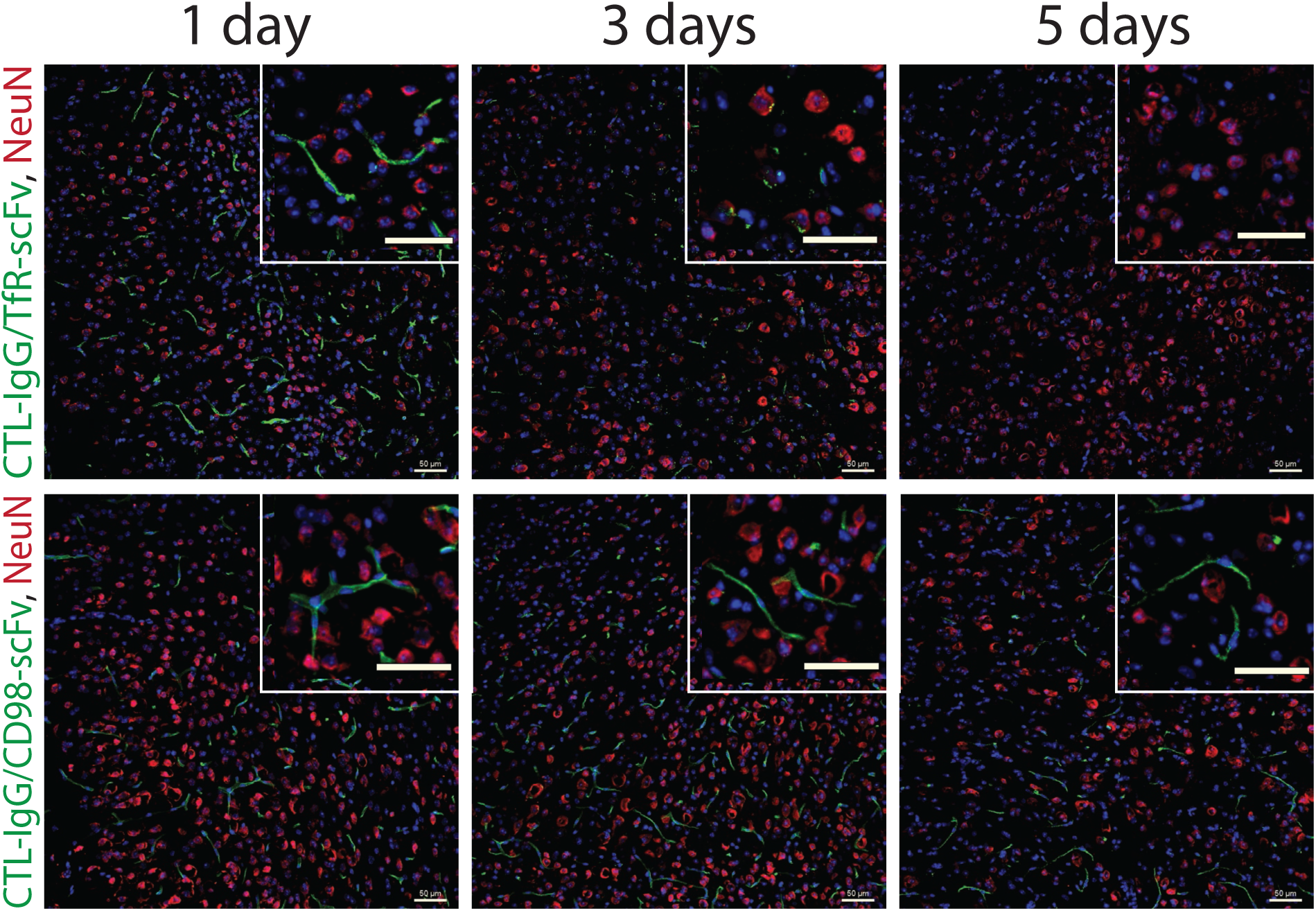
Brain distribution of CD98hc and TfR-1 shuttles delivering a non-targeted IgG. Brains were harvested at 1, 3 and 5 d after injection (3.6 mg/kg, 20 nmol/kg) of fluorescently labeled shuttles and sectioned for immunostaining. The injected shuttles (green) are shown relative to NeuN+ neurons (red) and nuclei (blue). Scale bars are 50 µm.

### Brain delivery of cell-specific and agonist IgGs

We next sought to evaluate the ability of the IgGs shuttled into the brain via CD98hc to engage targets in the brain parenchyma. To enable testing in wild-type mice, we searched for highly validated monoclonal antibodies that target endogenous, cell-surface antigens on different types of mouse brain cells. We identified two candidates, one specific for neurons (clone M6)^30^ and the other specific for astrocytes and oligodendrocytes (clone M2)^31, 32^, and reformatted the IgGs as 2×1 CD98hc shuttles.

For the M2/CD98hc shuttle specific for astrocytes, we observed little co-localization between the shuttle, which appeared to largely localize with blood vessels, and a marker of astrocytes (glial fibrillary acidic protein, GFAP), after one day (**Fig. 5**). However, three days after administration, there was overlap between M2/CD98hc shuttle and GFAP staining, and by five days we observed strong colocalization of the two fluorescent signals. Moreover, a 3 mg/kg dose of non-shuttled M2 IgG did not produce detectable fluorescent signal in the brain (**Fig. S11**).

**Figure 5.**
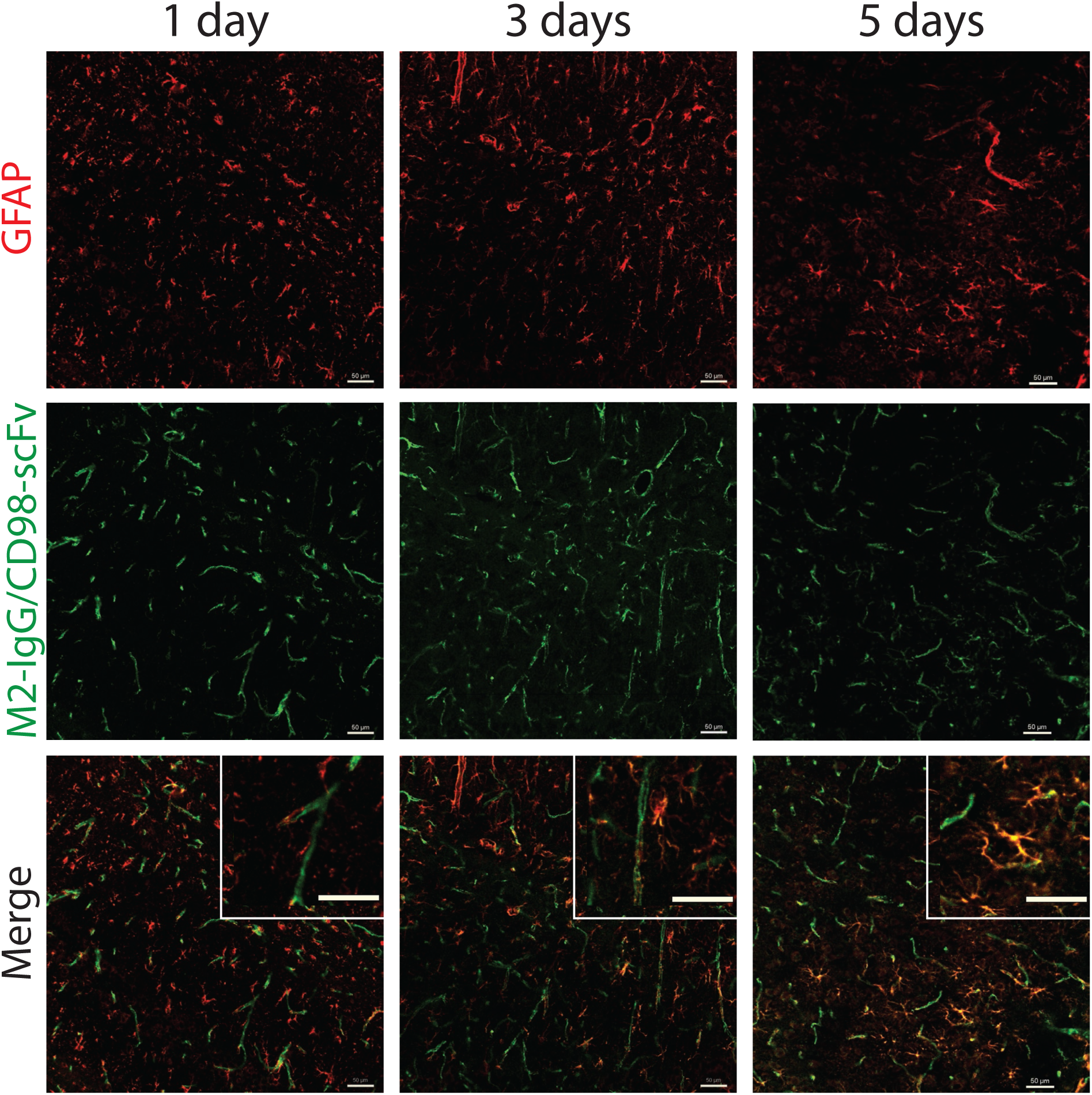
Brain distribution of a CD98hc-shuttled IgG specific for astrocytes. An astrocyte-specific IgG (M2), delivered via a CD98hc shuttle, was labeled with Alexa Fluor-647 and injected into wild-type mice (3.6 mg/kg, 20 nmol/kg). Brains were harvested after 1, 3 and 5 d, and sectioned for immunostaining. Antibody distribution of the shuttle (green) is shown relative to that of GFAP+ astrocytes (red). Scale bars are 50 µm.

The M6/CD98hc shuttle that is specific for neurons led to significantly different behavior (**Fig. 6**). Like the CD98hc-shuttled M2 antibody, a pattern suggestive of blood vessel localization was seen one day after administration, with no overlap with neuronal (NeuN) staining. After three days, the M6/CD98hc shuttle was strongly colocalized with NeuN+ neurons and appeared in a punctate distribution suggestive of cellular internalization. Finally, after five days, fluorescent signal from the shuttle was undetectable (**Fig. S12**). As with the M2 IgG, injection of non-shuttled M6 IgG produced little detectable fluorescent signal in the brain (**Fig. S12**).

**Figure 6.**
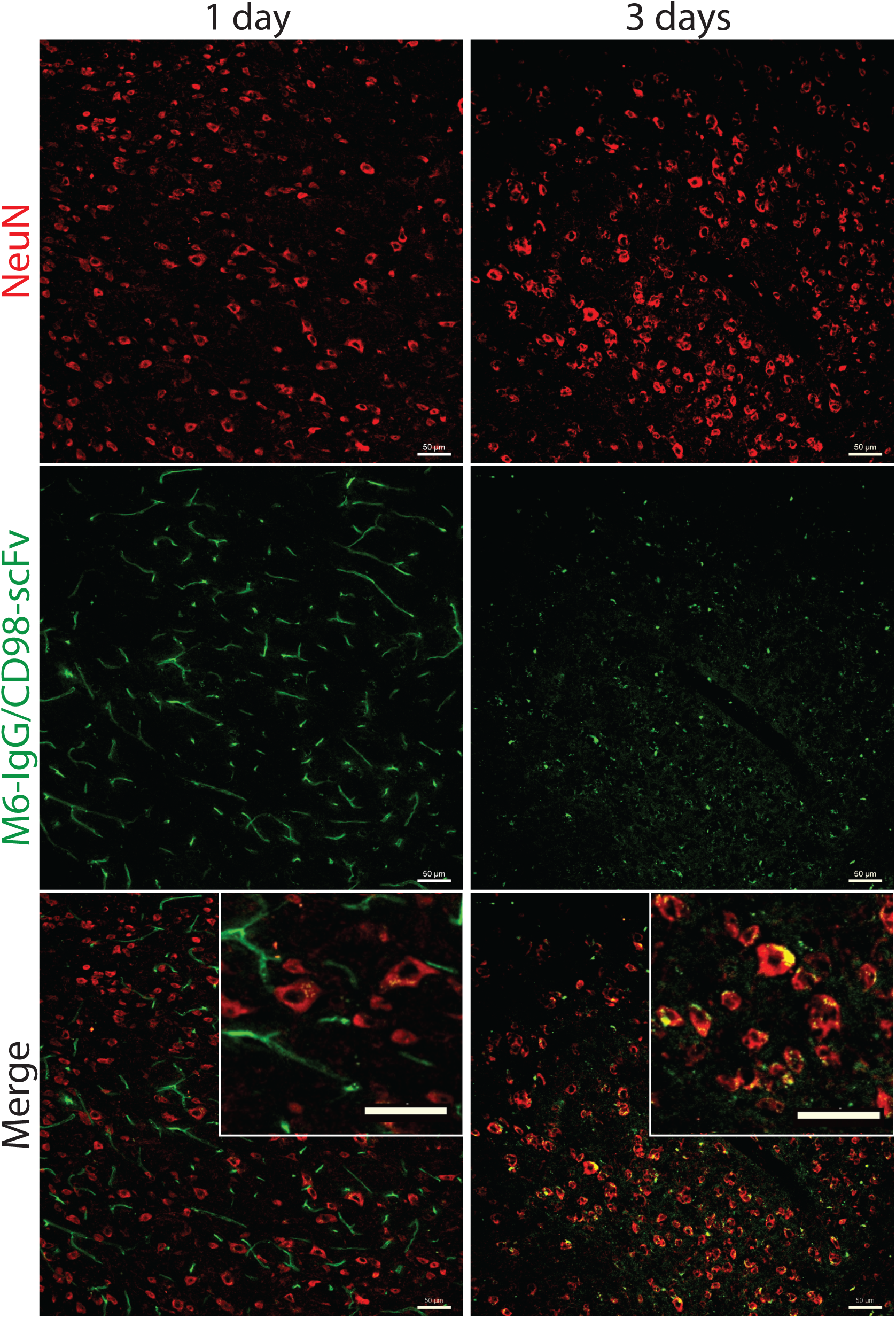
Brain distribution of a CD98hc-shuttled IgG specific for neurons. A neu-ron-specific IgG (M6), delivered via a CD98hc shuttle, was labeled with Alexa Fluor-647 and injected into wild-type mice (3.6 mg/kg, 20 nmol/kg). Brains were harvested after 1 and 3 d, and sectioned for immunostaining. Antibody distribution of the shuttle (green) is shown relative to that of NeuN+ neurons (red). Scale bars are 50 µm.

Finally, we sought to demonstrate the ability of the CD98hc shuttle to deliver an agonist antibody capable of activating an endogenous receptor in the mouse brain. Therefore, we selected an existing agonist IgG specific for the TrkB receptor with a published amino acid sequence, namely 29D7^33^ (**Fig. S1**). We reformatted the TrkB IgG as a 2×1 CD98hc shuttle, and confirmed its ability to recognize and activate TrkB *in vitro* in a similar manner as the corresponding IgG agonist (**Fig. S13**).

Next, we intravenously administered the TrkB IgG and shuttle, and evaluated their target engagement (i.e., TrkB receptor activation) using brain immunostaining analysis (**Fig. 7**). Similar to the M2 and M6 shuttles, the TrkB/CD98hc shuttle was predominantly localized to blood vessels after one day, although some signal was observed within NeuN+ neurons. At this same time point, we observed staining for phospho-Akt (p-Akt), a marker of TrkB signaling (**Fig. 7A**). A similar pattern was seen after three days, at which point, most of the TrkB/CD98hc shuttle was internalized into NeuN+ neurons. After five days, the shuttle was barely detectable and p-Akt staining was strongly reduced. Similar to the p-Akt immunostaining results, we observed staining for phospho-ERK1/2 (p-ERK1/2), another key signaling molecule downstream of the TrkB receptor, at 1 and 3 days after injection with the TrkB/CD98hc shuttle (**Figs. 7B** and **S14**). In contrast, injection of non-shuttled TrkB IgG produced little p-Akt or p-ERK1/2 staining (**Figs. S15** and **S16**). These results collectively demonstrate that the CD98hc-shuttled TrkB antibody strongly and specifically activates TrkB in the mouse brain for multiple days.

**Figure 7.**
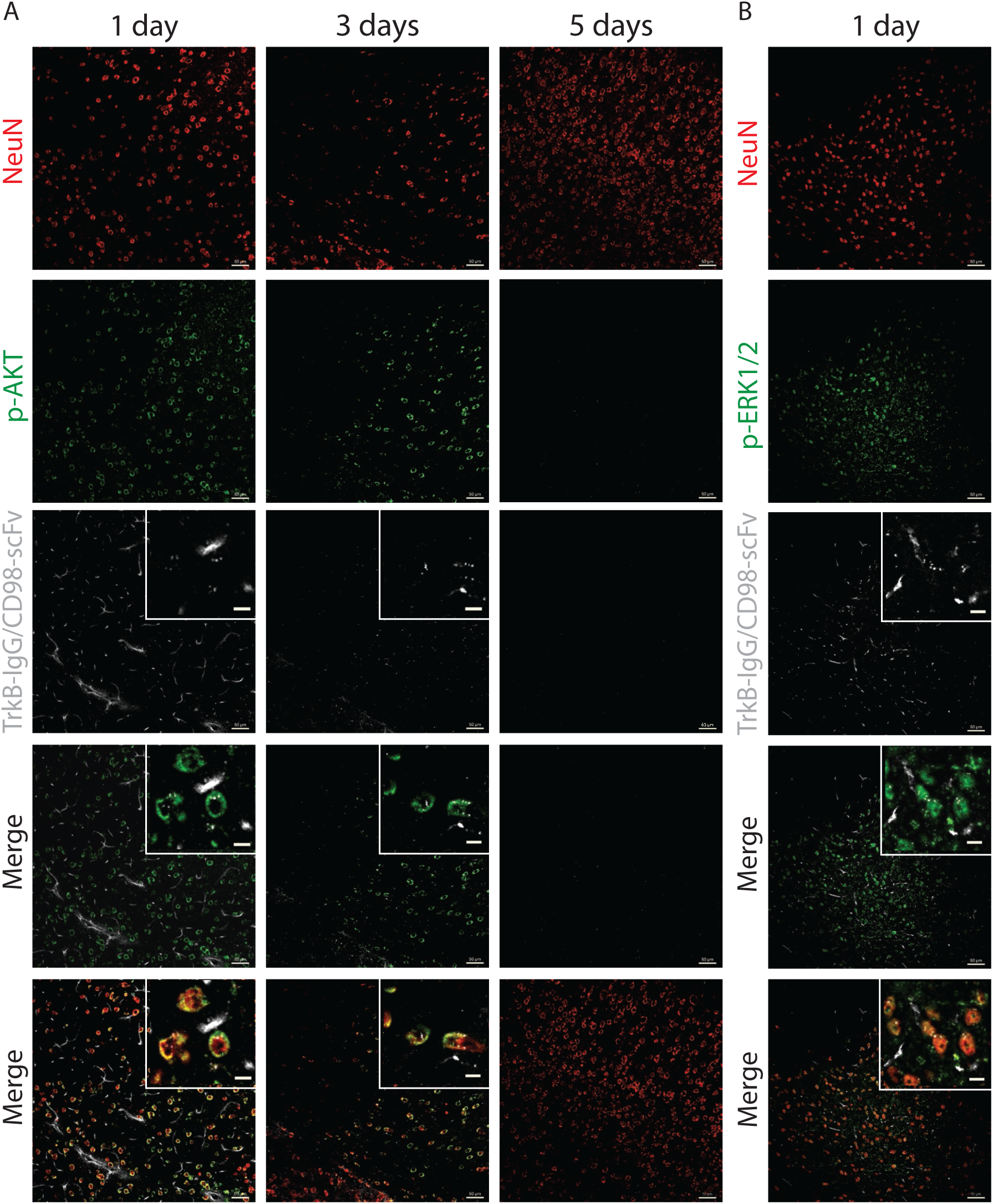
Brain distribution and TrkB activation of a CD98hc-shuttled agonist IgG. A TrkB agonist IgG, delivered via a CD98hc shuttle, was labeled with Alexa Fluor-647 and injected into wild-type mice (3.6 mg/kg, 20 nmol/kg). Brains were harvested after 1, 3 and 5 d, and sectioned for immunostaining. Antibody distribution of the shuttle (grey) is shown relative to that of NeuN+ neurons (red) and either p-AKT in (A) or p-ERK1/2 in (B) in green. Scale bars are 50 µm in the main images and 10 µm in the enlarged inset images.

## DISCUSSION

The high potency and extreme specificity of therapeutic IgGs have made them both the fastest growing drug class and essential tools for defining the molecular mechanisms of human disease. Despite their remarkable advantages, IgGs have had relatively little impact on the study or treatment of brain disorders, in large part, due to their inability to cross the BBB. Indeed, the recent failure of several, high-profile clinical trials has underscored the limitations of traditional antibody therapeutics for testing specific disease mechanisms within the brain^34, 35^. In the absence of a defined mechanism or kinetics of brain uptake or pharmacodynamic markers of target engagement, these trials have failed not only to achieve their intended clinical outcomes, but also to convincingly test the underlying therapeutic hypothesis^36^. With this in mind, we have focused on characterizing a BBB transport pathway with a defined molecular mechanism. Our current results represent detailed characterization of the kinetics of brain uptake, parenchymal retention, and target engagement of IgGs shuttled into the brain via CD98hc, with particular focus on the ways in which this transport pathway differs from that of TfR-1.

Perhaps the most remarkable and interesting finding of our work is the sustained brain concentration of IgGs shuttled across the BBB via CD98hc. Our PK data, confirmed by three different methods, indicate IgG concentrations more than half their peak value at one week after CD98hc shuttling, a time point at which TfR-1-shuttled IgG can no longer be detected in the brain. There are a number of plausible mechanisms which may explain the differences in delivery and retention of IgG shuttled by the two pathways. First, fluorescence imaging indicates profound differences in the localization of non-targeted IgG shuttled into the brain via CD98hc versus TfR-1. The former remains associated with, or at least in close proximity to, blood vessels for at least five days post-injection, whereas TfR-1 shuttled IgG appears to be internalized by multiple cell types in the brain parenchyma. These data suggest a relative lack of CD98hc engagement in the brain parenchyma, and conversely, robust TfR-1 engagement, leading to internalization, intercellular catabolism, and a faster reduction in brain levels of the TfR-1 shuttle^7, 9, 19, 20^. A second potential contributor is internalization and degradation of the cerebrovascular targets (i.e., TfR-1 and CD98hc) in the presence of saturating doses of 2×1 shuttle. This effect, which our *in vitro* analysis confirmed is far more pronounced for TfR-1 than CD98hc, would result in a steep decline in brain uptake and could explain both the earlier peak and the more rapid decline in brain concentration of the TfR-1 shuttle, relative to its CD98hc counterpart. Finally, there are potential differences in the polarization of CD98hc and TfR-1 on cerebrovascular endothelial cells^37–40^, which could impact and even bias bidirectional transport across the BBB and contribute to differences in brain accumulation and retention of the CD98hc and TfR-1 shuttles.

Another key aspect of our work is the 2×1 bispecific antibody format, which deserves further consideration. While there are many possible formats for bispecific antibodies^41^, including those that have been used as BBB shuttles^4, 8, 18, 20, 28, 42–49^, the 2×1 format reported previously for TfR-1 shuttles (IgG fused to one single-chain Fab)^8^ and used in this work (IgG fused to one scFv) is compelling for multiple reasons. First, the two Fab regions of the IgG portion of the shuttle are not modified in any way. This is a distinct advantage because it eliminates the potential for avidity changes and potential changes in Fab affinity and specificity for the shuttled IgG. In comparison, bispecific antibody formats, such as dual-variable-domain (DVD) shuttles, require changes to the Fab regions of the shuttled IgG, including the introduction of linkers between the outer and inner variable regions, which can impact the affinity of both the inner and outer variable domains^42, 50^. Most importantly, our results demonstrate straightforward conversion of existing IgGs (e.g., M2, M6, and 29D7) into 2×1 bispecific antibodies. There are presumably hundreds of similar antibodies with brain targets, developed for diagnostic or research applications, that have never been tested *in vivo* for their activity in the central nervous system, especially without the need for complex and highly invasive delivery methods.

Beyond its utility as a platform for delivering existing IgGs as reformatted brain shuttles, the other major advantage of the 2×1 format is its monovalent engagement of the BBB target^7, 48, 51^. Our data largely support the need for monovalent antibody engagement of TfR-1, which appears to reduce internalization and ultimately results in higher brain concentrations relative to the bivalent TfR-1 IgG. We observed mixed results for CD98hc IgG, which had higher peak brain concentrations than the 2×1 CD98hc shuttle at an intermediate dose (3 mg/kg for IgG and 3.6 mg for shuttle, 20 nmol/kg) and lower concentrations at a higher dose (9 mg/kg for IgG and 10.8 mg for shuttle, 60 nmol/kg). Given that the monovalent 2×1 CD98hc shuttle resulted in extended brain retention relative to its bivalent counterpart, monovalent engagement of CD98hc appears optimal for maximizing IgG brain levels and extended brain retention, assuming a sufficiently high dose is used. This observation will need to be investigated further given that the intrinsic CD98hc antibody affinity and epitope^52^, in addition to valency, are likely to impact the efficiency of IgG brain shuttling and retention.

It is also important to consider the implications of the unique biodistribution of the CD98hc and TfR-1 shuttles within the brain and between different organs. In the brain, we found that non-targeted CD98hc shuttles do not mediate appreciable cellular uptake in the brain parenchyma, unlike the corresponding TfR-1 shuttles. This potentially enables improved ability of CD98hc shuttles to specifically target different cell types in the brain. However, this increased specificity should be considered in the context of CD98hc-and TfR-1-mediated shuttling of antibodies to the brain versus other organs. In some cases, CD98hc shuttles led to increased non-brain organ uptake, including in the kidney and spleen, relative to TfR-1 shuttles. For certain applications, the reduced delivery of bispecific antibodies to certain non-brain organs may be an advantage for TfR-1 shuttles relative to CD98hc shuttles.

The interaction of CD98hc and TfR-1 shuttles with their cerebrovascular and parenchymal targets and their impact on brain uptake, retention, and target engagement are likely to have profound implications for the many exciting future directions of this work. First, the CD98hc shuttles, which result in limited engagement of CD98hc in the brain parenchyma, could be used in applications that seek to deliver IgGs to the brain without cellular internalization within the brain parenchyma. These applications could include delivery of IgGs that capture soluble factors such as Aβ in Alzheimer’s disease^53, 54^ and VEGF in glioblastoma^55, 56^. These tasks are more complex to perform using TfR-1 shuttles because of TfR-1– mediated shuttle internalization into cells within the brain parenchyma. Second, the ability of the CD98hc-shuttled M2 antibody to enter the brain parenchyma and remain stably associated with astrocytes could be exploited for the brain delivery of conjugated drugs. This may be particularly useful in applications where extracellular brain release of drugs is desired, such as antibiotics^57, 58^ or extended release of drugs into the brain parenchyma after brain localization to minimize systemic side effects, such as various types of toxins used for treating epilepsy^59^. Third, in contrast, the ability of the CD98hc-shuttled M6 antibody to specifically internalize into neurons could be utilized to deliver conjugated drugs that require intracellular delivery. For example, anti-sense oligonucleotides (ASOs) and small interfering RNAs (siRNAs) need intracellular delivery to evaluate the impact of altering protein expression on normal and disease phenotypes^60, 61^. This would be particularly significant because these agents are typically delivered to the brain by invasive methods (e.g., intrathecal or intracerebroventricular administration)^62–65^. Fourth, the ability of the CD98hc-shuttled TrkB antibody to activate both AKT and ERK1/2 for at least two days after a single intravenous dose, which is one of the longest periods of *in vivo* TrkB activation reported to date after a single dose^66–,68^, could be used to evaluate the therapeutic impact of activating TrkB in the context of several acute (e.g., traumatic brain injury and stroke) and chronic (e.g., neurodegenerative) diseases. This is significant because there is preclinical evidence that TrkB activation is neuroprotective^69–75^ and/or reduction of brain-derived neurotrophic factor (BDNF) or TrkB signaling is linked to neuronal dysfunction^76–80^. These and other exciting possibilities are expected to simplify fundamental and therapeutic studies of the delivery of antibodies and other agents to the brain.

## MATERIALS AND METHODS

### Recombinant antibodies

Antibody sequences are available in the supplementary section (**Fig. S1**). CTL IgG is an anti-phospho-tau IgG (PDB ID: 5DMG), which was generated by rat immunization using phosphorylated tau (pS422)^81^. CD98hc IgG is an anti-mouse CD98hc IgG generated by rat immunization using mouse bone marrow cells^21^. TfR-1 IgG (clone 8D3) is an anti-mouse TfR-1 IgG prepared by rat immunization using murine endothelial cells (t.end 1)^82^. M2 is an anti-mouse astrocyte surface antigen IgG produced by rat immunization using the particulate fraction of mouse cerebellar homogenates^31^. M6 is an anti-mouse neuronal membrane glycoprotein M6-a IgG generated by rat immunization using mouse cerebellar homogenate^30^. The variable regions of M2 and M6 hybridomas were sequenced after generating cDNA internally or by GenScript with permission from Dr. Carl Lagenaur, University of Pittsburgh, and University of Heidelberg. 29D7 (TrkB-IgG) is an anti-TrkB IgG produced via mouse immunization using the extracellular domains of human and mouse TrkB^83^.

### Cloning and production of IgGs and bispecific shuttles

Expression vector pTT5 and mammalian cell line (HEK293-6E) were licensed from the National Research Council of Canada (NRC). A bispecific shuttle was generated by first introducing ‘knob-into-hole’ mutations^22, 23^ in the Fc region of human IgG1. Next, genes of either TfR-1 scFv or CD98 scFv were genetically fused via a glycine-serine linker (G_4_S)_3_ to the C-terminus of the ‘knob’ chain. The expression vector was digested with appropriate restriction enzymes (New England Biolabs) following the manufacturer’s protocol, and subsequently treated with calf intestinal alkaline phosphatase (New England Biolabs, M0525L). The digested vector was separated by gel electrophoresis (1% agarose). Appropriately sized DNA was excised and purified (Qiagen, 28704). Digested inserts and vectors were ligated with T4 ligase (New England Biolabs, M0202L) and transformed via heat shock into DH5α competent cells. Cells were incubated in the presence of LB media (antibiotic free) for 1 h at 37 °C on an orbital shaker (200 rpm), and then plated on LB plates supplemented with ampicillin (100 µg/mL) overnight at 37 °C. Single colonies were picked, grown in ampicillin-supplemented LB media overnight, miniprepped (Qiagen, 27106), and sequenced by Sanger sequencing.

Expression of antibodies was performed by transfecting 25 mL of mammalian cell culture (2×10^6xref>^ cells/mL) with appropriate plasmids. In each conical tube, 15 µg of total DNA (equal parts knob, hole, and light chains) was mixed with 45 µL of 40 kDa polyethylenimine (1 mg/mL) and 3 mL of F17 media (Thermo Fisher, A1383501) in a sterile hood for 15 min. Next, the mixture was added to the cells and then transferred back to the incubator (37 °C, 5% CO_2_). After 24 h, 0.75 mL of yeastolate (%20 w/w) (Thermo Fisher, B92804) was added to each tube of transfected cells. The cells were harvested 6 d after transfection and centrifuged at 3500 rpm for 40 min.

The supernatant was filtered (Thermo Fisher, 166-0045) and passed through a pre-packed Protein A agarose (Thermo Fisher, 20333) column. After washing the column with 50 mL of PBS, 20 mL of glycine buffer (pH 3.0) was added to the column to elute the protein. Elution fractions were collected, and immediately buffer exchanged into 20 mM Acetate (pH 5.0) using desalting columns (Thermo Fisher, 89891) and stored at −80 °C. The absorbance (A_280_) was measured via NanoDrop to determine antibody concentration. Antibodies were run on SDS-PAGE gels as well as analyzed via size exclusion chromatography (SEC) (Superdex 200 Increase 10/300 GL, Cytvia #28990944). Antibodies that were <90% monomer after Protein A purification, as determined by SEC, underwent a second-step purification via fraction collection on SEC until desired purity was achieved.

### Generation of mouse TfR and CD98hc expressing cell lines

Full-length coding sequences for murine TfR-1 (CD71, NCBI reference sequence: NP_035768.1) and CD98hc (NCBI reference sequence: NP_001154885.1) were cloned into the pcDNA 3.1 (+) plasmid between restriction enzyme sites NheI/XbaI and HindIII/EcoRI, respectively. For CD98hc, a FLAG tag was fused to the intracellular N-terminal domain. Plasmids were transfected into the REN cell line using Lipofectamine 2000 per manufacturer protocol^84^. Cells were selected in media containing 200 µg/mL of Geneticin (Thermo Fisher Scientific) and flow sorted using a MoFlo Astrios cell sorter (Beckman Coulter, Brea, CA) based on binding of TfR1 and CD98hc IgG, respectively.

### Flow cytometry binding analysis of TfR-1 and CD98hc shuttles

TfR-expressing (REN-TfR) and CD98-expressing (REN-CD98) cell lines were grown in RPMI media with 10% FBS, 1% Antibiotic-Antimycotic (“anti-anti”), and 200 μg/mL Geneticin until confluent. Cells were washed with PBS, detached from the flask with 0.25% Trypsin-EDTA, and resuspended in fresh media at a density of 10^6^ cells/mL. Cells were then seeded on a 96-well plate at a density of 10^5^ cells per well. Serial dilutions of antibodies were prepared with cold PBS supplemented with 0.1% BSA (PBSB). Seeded cells were treated with serial dilutions of the antibodies for 1 h at 4 °C on an orbital shaker (200 rpm). After treatment, the plate was centrifuged at 2500 rpm for 4 min and washed with PBSB twice. The cells were then incubated with an anti-human Fc detection antibody AF-647 (1:500; Jackson, 109-605-098) for 4 min on ice. After incubation, the plate was centrifuged at 2500 rpm for 4 min and washed with PBSB twice. The cells were resuspended with 200 μL of PBSB and analyzed on a flow cytometer (Bio-Rad, ZE5 cell analyzer).

### Flow cytometry analysis of IgG and bispecific affinity for soluble CD98hc and TfR-1 ectodomains

Affinities of IgGs and bispecifics were measured by flow cytometry after immobilizing antibodies on the surface of Protein A Dynabeads. Protein A Dynabeads (Invitrogen, 10002D) were washed with 1x PBS supplemented with 0.1% BSA (PBSB) and incubated with antibodies (85 µL at 15 µg/mL) overnight at 4°C with mild agitation. The beads were then washed with PBSB and incubated with biotinylated CD98hc and TfR-1 ectodomains at a range of concentrations (1250, 250, 50, 10, 2, 0.4, 0.08 and 0.016 nM) such that antigen was present in 10-fold molar excess of immobilized antibodies at 4 °C with mild agitation for approximately 3 h.

CD98hc IgG and CD98hc bispecific antibodies were incubated with mouse CD98hc ectodomain. The CD98hc ectodomain (NP_001154885 residues 139-565 with an N-terminal 6x His tag) was expressed in HEK293-6E cells, as previously described for the production of IgGs and mAbs,^85, 86^ and purified using immobilized metal-affinity chromatography (Qiagen, 30230) and size-exclusion chromatography. Briefly, supernatant containing mouse CD98hc ectodomain was incubated with nickel beads overnight at 4 °C, the beads were washed with 1x PBS and 50 mM imidazole, and ectodomain was eluted from the beads with 500 mM imidazole. The CD98hc ectodomain was then buffer exchanged into 1x PBS, and monomeric CD98hc was further purified by size-exclusion chromatography. Purified CD98hc was aliquoted and stored at −80 °C until use.

TfR IgG and TfR bispecific antibodies were incubated with mouse TfR (ACRO Biosystems, TFR-M524b). The beads were then washed with PBSB and incubated with a 1:1000 dilution of streptavidin Alexa Fluor 647 (Invitrogen, S32357) and 1:1000 dilution of goat anti-human Fc F(‘ab)_2_ Alexa Fluor 488 (Jackson ImmunoResearch, 109-546-170) on ice for 4 min. Next, the beads were washed once with PBSB and resuspended in PBSB for analysis by flow cytometry using a Bio-Rad ZE5 cell analyzer. The mean fluorescent intensities were recorded and analyzed. Results are reported after subtraction of background signal observed in the absence of antigen (0 nM), subtraction of the lowest binding signal observed (baseline of 0), and normalization to the highest binding signal observed.

### Endothelial surface level analysis of TfR-1 and CD98hc

Mouse brain endothelial cells (bEnd.3, ATCC, CRL-2299) were plated on a 6-well plate at a density of 3×10^5^ cells/well and grown overnight in 2 mL of growth media (DMEM with 10% FBS and 1% anti-anti). Cells were then treated overnight with either 100 nM IgG (TfR or CD98) or 200 nM shuttle (CTL-IgG/TfR-or CD98-scFv) added to the growth media. The next day, media was removed and cells were washed with PBS then treated with 0.25% Trypsin-EDTA for 3-5 min until cells became detached. Cells were then resuspended in 1mL of growth media and transferred to microcentrifuge tubes. Cells pre-treated with anti-mouse TfR-1 antibodies (human constant regions) were incubated with 50 nM of a second TfR-1 antibody for detection (R17, rat anti-mouse antibody; BioLegend, 113802). Cells pre-treated with anti-mouse CD98hc antibodies (human constant regions) were incubated with 10 nM of a second CD98hc antibody for detection (RL388, rat anti-mouse antibody; BioLegend, 128202). Non-treated cells were also incubated with the TfR-1 (R17) and CD98hc (RL388) detection antibodies to determine baseline levels of TfR-1 and CD98hc without antibody-mediated internalization. The incubations with the detection antibodies (R17 and RL388) lasted for 1 h and were performed while rotating at 4 °C. Cells were then twice spun down at 2500 rpm for 4 min and resuspended in PBSB. Following the second wash, cells were incubated for 4 min, on ice, with a secondary (anti-rat) detection antibody labeled with Alexa Fluor-647 (Biolegend, 407511) at a 1:500 dilution. Samples were again twice spun down and washed with PBSB. Cells were resuspended in a final volume of 100-200 μL PBSB and analyzed on a flow cytometer (Bio-Rad, ZE5 cell analyzer).

### Animal use and protocol

Animal studies were conducted following guidance for the care and use of laboratory animals as adopted by the NIH, under protocols PRO00010991 and PRO00009238 approved by the institutional animal care and use committee (IACUC) of the University of Michigan. C57BL/6J male mice (Jackson Laboratory, 000664), aged 8-16 weeks, were used for all animal experiments.

### Radiolabeling

Proteins were directly radioiodinated with [^125^I]NaI (Perkin Elmer) using Pierce iodination reagent (Thermo Fisher, 28601) and purified using Zeba desalting columns. Radiochemical purity was assessed via thin layer chromatography (TLC) performed using aluminum TLC silica gel 60 F254 plates (Millipore Sigma, 105554) and a 75%:25% mixture of methanol and 1 M sodium acetate (pH 6.8) as a mobile phase, as previously reported^87^. Radiolabeling efficiency was typically >75% and proteins were only used for radiotracing if free ^125^I was <5% following purification.

### Pharmacokinetic analysis

For all radiotracing experiments, a tracer dose (∼1 μg) of ^125^I-labeled IgG or bispecific shuttle was added to the appropriate dose of unlabeled protein to achieve approximately 10^6^ cpm on a Wizard2 2470 gamma counter. Prior to injection, mice were weighed, and doses were sterile-filtered and prepared in sterile PBS to a final volume of 110 μL. Mice were sedated with isoflurane (Henry Schein, 66794-017-25) and injected intravenously via the retroorbital plexus. In some experiments, mice were re-sedated briefly (<5 min) at intermediate time points for blood collection using heparin-coated capillary tubes (Thermo Fisher, 22-3652566). At the terminal time point, mice were re-sedated and transcardially perfused with ice-cold PBS. Blood and organs of interest were harvested and weighed prior to gamma counting.

### Antibody brain parenchyma concentration analysis

^125^I-labeled antibody doses were prepared as described above at concentrations of 3 mg/kg for IgGs (20 nmol/kg) and 3.6 mg/kg for shuttles (20 nmol/kg). One hour after injection, mice were anesthetized with isoflurane and transcardially perfused with ice-cold PBS with 10% trypan blue (Gibco, 15250061). Trypan blue was used to visualize brain vasculature after homogenization. Brains were harvested and homogenized in 1 mL Krebs buffer (Thermo Fisher, J67591.AP) using a TissueLyser LT (Qiagen, 85600) for 5 min at 40 Hz. The resulting homogenate was then added to 9 mL of 18% dextran (Acros, 406271000) in a 15 mL conical tube, shaken vigorously, and then centrifuged at 4300x*g* in a swinging bucket rotor for 45 min. Capillary depletion was confirmed through visualization of a pellet containing the trypan blue stained vessels while brain parenchyma remained layered above the dextran. Fractions of 1 mL were drawn from the top of the tube and separated into glass vials. The vials were analyzed on the gamma counter and the resulting counts were used to determine the percent of ^125^I-labeled antibodies in the capillaries and parenchyma.

### ELISA for the determination of brain antibody concentration

Immulon 4HBX flat bottom plates (Thermo Fisher, 3855) were coated with anti-human polyvalent immunoglobulins (Sigma, I1761) in a 1:1000 dilution and stored at 4 °C overnight. Harvested brains were weighed and 1 mL of RIPA buffer, including 10 µL of 100X Halt Protease Inhibitors (Thermo Fisher, 1860932), was added to each brain sample. Brains were homogenized using a Qiagen TissueLyser LT and the resulting homogenate incubated in RIPA buffer overnight at 4 °C. The homogenate was then centrifuged at 14,000x*g* for 20 min and the supernatant was recovered for ELISA analysis. ELISA plates were washed 4x with 0.5% PBST, blocked for 1 h with 5% goat serum, and then supernatant was added to ELISA plates in serial dilution (100 µL) prepared in 5% goat serum. The corresponding antibody was also prepared in RIPA buffer with protease inhibitors to be used as a standard curve. Samples incubated for 2 h and were then washed 4x with 0.5% PBST. 100 µL of anti-human IgG (Fc specific)-peroxidase antibody produced in goat (Sigma, A0170) was added to each sample in a 1:40,000 dilution of PBS and incubated for 1 h. Plates were washed 4x with 0.5% PBST, incubated for 15 min in TMB Substrate Solution (Thermo Fisher, N301), and quenched with 0.18 M H_2_SO_4_. The plates were then analyzed for their absorbance at 450 nm, and concentrations were calculated based on the standard curve and the dilution factor calculated from the initial weight of the brain.

### Fluorescence imaging of antibody organ distribution

Antibodies were first labeled with Alexa Fluor-647 NHS Ester (Thermo Fisher, A20006) following manufacturer’s protocol to achieve a degree of labeling of ∼1.0. Briefly, 200 µL of antibody (1 mg/mL) in PBS was mixed with 20 µL of sodium bicarbonate (1 M) and 0.4 µL of dye (10 mg/mL). Purification was performed by centrifuging (1 min, 12000 rpm) the labeled sample in a 30 kDa molecular weight cutoff centrifugal filter (Millipore Sigma, UFC503096) and washing the retained labeled antibody with 200 µL of PBS three times. The absorbances (280 and 650 nm) were measured on a NanoDrop to calculate the degree of labeling.

C57BL/6J male mice (8-16 weeks old) were intravenously injected with either labeled IgGs (3 mg/kg) or bispecific shuttles (3.6 mg/kg). After 7 days, the mice were sacrificed and transcardial perfusion was performed with 25 mL of ice-cold PBS. Organs including the liver, lung, kidneys, heart, spleen, and brain were removed and transferred to a 6-well plate. *Ex vivo* imaging of the organs was performed on a fluorescence imager (Perkin Elmer, IVIS Lumina S5).

### Brain sectioning and immunofluorescence analysis

After transcardial perfusion with ice-cold PBS, the mouse brains were removed and submerged in 4% paraformaldehyde (Thermo Fisher, J19943.K2) overnight at 4 °C for fixation. The brains were then transferred to a 30% sucrose solution for 24 h at 4 °C. The whole brains were first divided into two sagittal hemispheres. Next, the right hemispheres were divided coronally in the middle with a razor, placed in a cryomold filled with OTC and transferred to a −80°C freezer. Sections (20 µm) were collected by using a cryostat (Leica, NX50), which were then transferred onto glass slides (Thermo Fisher, 1255015), and left to dry overnight at room temperature.

Immunostaining was performed using standard protocols^88, 89^. A barrier was first drawn around the brain section using a hydrophobic pen (Vector laboratories, H-4000). The section was treated with cold methanol for 5 min and washed with PBS. Antigen retrieval was performed by heating the slides in 10 mM sodium citrate buffer pH 6.0 (Thermo Fisher, S279-3) for 4 min in a microwave and left to cool down for 10 min at room temperature. The sections were washed with PBS and treated with 0.5% Triton-X 100 (Alfa Aesar, A16046) for 10 min. The sections were then blocked with PBS supplemented with 5% goat serum (Novus Biologicals, S13150NOV) and 0.05% Triton-X 100 for 1 h. The sections were washed with PBS and then incubated with primary antibodies (**Table S2**), prepared in blocking solution using the manufacturer’s suggested dilution, overnight at 4°C. Next, the sections were washed with PBS and incubated with appropriate secondary detection antibodies (**Table S2**), following manufacturer’s suggested dilutions, for 1 h at room temperature. The sections were washed with PBS and counter stained with DAPI (1:1000) for 4 min. The sections were washed with PBS twice, preserved with an antifade reagent (Thermo Fisher, P36930), and protected with a coverslip (Thermo Fisher, 12543D). Immunofluorescence imaging was performed using a confocal microscope (Nikon, A1si confocal).

### *In vitro* TrkB affinity analysis

Mouse TrkB-expressing (REN-mTrkB) cells were grown in RPMI media with 10% FBS, 1% anti-anti, and 200 µg/mL Geneticin until confluent. Cells were washed with PBS, detached from the flask with 0.25% trypsin, and resuspended in 2% milk in PBSB to a density of 10^6^ cells/mL. Cells were then seeded on a 96-well plate at a density of 10^5^ cells/well. Serial dilutions of antibodies (TrkB-IgG and TrkB-IgG/CD98-scFv) were prepared with cold PBSB. Seeded cells were treated with serial dilutions of the antibodies for 1 h at 4 °C on an orbital shaker (200 rpm). After treatment, the plate was centrifuged at 2500 rpm for 4 min and washed with PBSB twice. The cells were then incubated with an anti-human Fc detection antibody AF-647 (1:500; Jackson, 109-605-098) for 4 min on ice. After incubation, the plate was centrifuged at 2500 rpm for 4 min and washed with PBSB twice. The cells were resuspended with 200 µL of PBSB and analyzed on a flow cytometer (Bio-Rad, ZE5 cell analyzer).

### Western blot analysis

REN cells expressing mouse TrkB (REN-mTrkB) were seeded into 6-well plates (3×10^5^ cells/well) and grown to confluence in RPMI media with 10% FBS, 1% anti-anti, and 200 μg/mL Geneticin. Once confluent, media was removed, and cells were treated with 1 nM TrkB IgG, 1 nM TrkB shuttle, 10 nM TrkB IgG, and 10 nM TrkB shuttle in RPMI media. Additionally, cells were treated with 1 nM BDNF as a positive control and 10 nM of CTL-IgG as a negative control. Cells were incubated at 37 °C for 15 min. Media was removed and cells were washed with PBS twice. 200 μL of RIPA buffer with phosphatase and protease inhibitors (Thermo Fisher, P178440) and 2% SDS was added to each well and incubated on ice for 30 min. Cells were then scraped from the wells and transferred to a microcentrifuge tube on ice. Samples were boiled at 100 °C for 5 min and centrifuged at max speed (∼21000×g) for 5 min. Protein concentration of each sample was determined using a Pierce™ BCA Protein Assay Kit (Thermo Fisher, 23227).

Next, the samples were combined with 4x loading buffer (40 mM Tris-HCl, 0.04% (w/v) ý-mercaptoethanol, 40% glycerol, 4 mM EDTA, 0.2 mg/ml bromophenol blue and pH 6.8). 15 μg of each sample was loaded onto a 4-12% Bis-Tris SDS-PAGE gel (Thermo Fisher, WG1401A). Two μL of MagicMark XP Western Protein Standard (Thermo Fisher, LC5602) was loaded with each gel. Proteins were transferred from the SDS-PAGE gels to nitrocellulose membranes (Thermo Fisher, IB23001) using the iBlot 2 Dry Transfer System (Thermo Fisher, IB21001) at 20 V for 1 min followed by 23 V for 4 min and 25 V for 2 min. The blots were then subsequently blocked for 1 h with 5% milk in TBS with 0.5% Tween-20 (TBS-T). The blots were next washed three times for 5 min each with TBS-T. Primary antibodies were diluted in 5% milk in TBS-T and incubated with the membrane rocking overnight at 4 °C. The blots were then washed and incubated for 1 h in goat anti-rabbit HRP conjugated secondary antibody (Cell Signaling Technology, 7074) diluted 1:3000 in TBS-T. The blots were washed three times for 5 min each with TBS-T then incubated in SignalFire ECL Reagent (Cell Signaling Technology, 6883) for 1 min and imaged using a BioRad ChemiDoc imager.

### Statistical analysis

All statistical analyses were performed in GraphPad Prism 9.0 using either a two-tailed unpaired t-test (α=0.05) with Welch’s correction or a one-way ANOVA with multiple comparisons using Tukey’s test for significance (α=0.05). Unless otherwise noted, all data presented as the mean ± one S.D. with *n* = 3-5 replicates.

## Data availability

Source data are provided with the paper and are available from the authors upon reasonable request.

## ACKNOWLEDGEMENTS

We thank members of the Tessier and Greineder labs for their helpful suggestions, Gabriel Corfas for assistance in analyzing the TrkB antibodies, Greg Thurber for suggesting brain delivery of the M2 and M6 antibodies, Richard Keep and Anuska Andjelkovic-Zochowska for advice on brain delivery and immunostaining, and Carl Lagenaur for generating the M2 and M6 hydridomas and providing permission to sequence the corresponding antibodies. This work was supported by the Massey Foundation at the University of Michigan (C.F.G. and P.M.T), National Institutes of Health (R01AG080016 to P.M.T. and C.F.G., RF1AG059723 and R35GM136300 to P.M.T., K08HL130430 to C.F.G., and T32 NS007222 and F32 AG079576 fellowships to M.J.L.), National Science Foundation (CBET 1813963, 1605266 and 1804313 to P.M.T.), BrightFocus Foundation (A2017395S to P.M.T. and A2022050S to P.M.T. and C.F.G), Coins for Alzheimer’s Research Trust (John Trojanowski Memorial Award to P.M.T. and C.F.G.), and the Albert M. Mattocks Chair (to P.M.T).

## AUTHOR CONTRIBUTIONS

G.P. and P.T. conceived the 2×1 CD98hc shuttle. P.T. and C.G. co-led the development of the 2×1 CD98hc shuttle. G.P., M.L. and J.Z. cloned the antibodies, and G.P. and M.L. produced and purified the antibodies. G.P., M.L., L.B. and C.G. performed pharmacokinetic analysis and P.T. provided significant input. G.P. and Y.K. performed brain immunostaining. C.G. and B.Z. prepared cell lines for flow cytometry and western blotting analysis. G.P., L.B. and J.Z. performed flow cytometry characterization of antibody binding. M.L. and L.B. performed western blotting. P.T., C.G., G.P., L.B. and M.L. wrote the manuscript with feedback from co-authors.

