## Supplemental Figures 1-16 and Tables 1-2 for "Bispecific antibody shuttles targeting CD98hc mediate efficient and long-lived brain delivery of IgGs"

A

| Antibody | $V_H$ | $V_L$ |
|---|---|---|
| CTL-IgG | QSVEESGGRLVTPGTPLTLTCTVSGFSLSSNAINWVRQAPGKGLEWIGYIAVSGN<br>TYYASWAKGRFTISKASTTVDLKMTSPTAEDTGTYFCGKSNIWGPGTLVTVSL | QVLTQTTSPVSAAVGSTVTISCQSSQSVRTNKLAWFQQKPGQPPKRLIYSASTLDF<br>GVPSRFSASGSGTQFTLTISDVQCDDAATYYCLGYFDCSIADCVAFGGGTEVVVK |
| TfR-IgG | EVQLVESGGGLVQPGNSLTLSCVASGFTFSNYGMHWIRQAPKKGLEWIAMIYYD<br>SSKMNYADTVKGRFTISRDNSKNTLYLEMNSLRSEDTAMYYCAVPTSHYVVDVW<br>GQGVSVTVSS | DIQMTQSPASLSASLEEIVTITCQASQDIGNWLAWYQQKPGKSPQLLIYGATSLAD<br>GVPSRFSGSRSGTQFSLKISRVQVEDIGIYYCLQAYNTPWTFGGGTKLELK |
| CD98-IgG | EVQLVESGGGLVQPGRSLKLSCAASGFTFSDYYMAWVRQAPKKGLEWVASISYE<br>GSSIYYGDSVKGRVTISRDNAKSTLYLQMNSLRSEDTATYYCARRGYYGYKPFDY<br>WGQGVMVTVSS | DIQMTQSPASLSASVGETVTIECRASEDIYNGLAWYQQKPGKSPQLLIYNIDNLHTG<br>VPSRFSGSGSGTQYSLKINSLQSEDVASYFCQQYYTYAYTFGAGTKLELK |
| TrkB-IgG | EVQLVESGGGLVQPGGSLRLSCAASGYSFTAYFMNWVRQAPGKGLEWVARINP<br>NNGDTFYTQKFKGRFTISRDNAKNSLYLQMNSLRAEDTAVYYCARRDYFGAMDY<br>WGQGTLVTVSS | DIQMTQSPSSLSASVGDRVTITCRASQTISNNLHWYQQKPGKAPKLLIKSASLAISG<br>VPSRFSGSGSGTDFTLTISSLQPEDFATYYCQQSNSWPNTFGGGTKVEIK |

B

| Chain | Sequence |
|---|---|
| Heavy chain* #1<br>(Knob, CD98hc) | ASTKGPSVFPLAPSSKSTSGGTAALGCLVKDYFPEPVTVSWNSGALTSGVHTFPAVLQSSGLYSLSSVVTVPSSSLGTQTYICNVNHKPSNTKVDKKVEP<br>KSCDKTHTCPPCPAPEAAGGPSVFLFPPKPKDTLMISRTPEVTCVVVDVSHEDPEVKFNWYVDGVEVHNAKTKPREEQYNSTYRVVSVLTVLHQDWLN<br>GKEYKCKVSNKALGAPIEKTISKAKGQPREPQVYTLPPCRDELTKNQVSLWCLVKGFYPSDIAVEWESNGQPENNYKTTPPVLDSDGSFFLYSKLTVDK<br>SRWQQGNVFSCSVMHEALHNHYTQKSLSLSPGKGGGGSGGGGSGGGGSDIQMTQSPASLSASVGETVTIECRASEDIYNGLAWYQQKPGKSPQLLI<br>YNIDNLHTGVPSRFSGSGSGTQYSLKINSLQSEDVASYFCQQYYTYAYTFGAGTKLELKRGGGGSGGGGSGGGGSGGGGSEVQLVESGGGLVQPG<br>RSLKLSCAASGFTFSDYYMAWVRQAPKKGLEWVASISYEGSSIYYGDSVKGRVTISRDNAKSTLYLQMNSLRSEDTATYYCARRGYYGYKPFDYWGQG<br>VMVTVSS |
| Heavy chain* #2<br>(Knob, TfR-1) | ASTKGPSVFPLAPSSKSTSGGTAALGCLVKDYFPEPVTVSWNSGALTSGVHTFPAVLQSSGLYSLSSVVTVPSSSLGTQTYICNVNHKPSNTKVDKKVEP<br>KSCDKTHTCPPCPAPEAAGGPSVFLFPPKPKDTLMISRTPEVTCVVVDVSHEDPEVKFNWYVDGVEVHNAKTKPREEQYNSTYRVVSVLTVLHQDWLN<br>GKEYKCKVSNKALGAPIEKTISKAKGQPREPQVYTLPPCRDELTKNQVSLWCLVKGFYPSDIAVEWESNGQPENNYKTTPPVLDSDGSFFLYSKLTVDK<br>SRWQQGNVFSCSVMHEALHNHYTQKSLSLSPGKGGGGSGGGGSGGGGSDIQMTQSPASLSASLEEIVTITCQASQDIGNWLAWYQQKPGKSPQLLI<br>YGATSLADGVPSRFSGSRSGTQFSLKISRVQVEDIGIYYCLQAYNTPWTFGGGTKLELKRGGGGSGGGGSGGGGSGGGGSEVQLVESGGGLVQPG<br>NSLTLSCVASGFTFSNYGMHWIRQAPKKGLEWIAMIYYDSSKMNYADTVKGRFTISRDNSKNTLYLEMNSLRSEDTAMYYCAVPTSHYVVDVWGQGVS<br>VTVSS |
| Heavy chain* #3<br>(Hole) | ASTKGPSVFPLAPSSKSTSGGTAALGCLVKDYFPEPVTVSWNSGALTSGVHTFPAVLQSSGLYSLSSVVTVPSSSLGTQTYICNVNHKPSNTKVDKKVEP<br>KSCDKTHTCPPCPAPEAAGGPSVFLFPPKPKDTLMISRTPEVTCVVVDVSHEDPEVKFNWYVDGVEVHNAKTKPREEQYNSTYRVVSVLTVLHQDWLN<br>GKEYKCKVSNKALGAPIEKTISKAKGQPREPQVCTLPPSRDELTKNQVSLSCAVKGFYPSDIAVEWESNGQPENNYKTTPPVLDSDGSFFLVSKLTVDK<br>SRWQQGNVFSCSVMHEALHNHYTQKSLSLSPGK |
| Heavy chain* #4<br>(Control) | ASTKGPSVFPLAPSSKSTSGGTAALGCLVKDYFPEPVTVSWNSGALTSGVHTFPAVLQSSGLYSLSSVVTVPSSSLGTQTYICNVNHKPSNTKVDKKVEP<br>KSCDKTHTCPPCPAPEAAGGPSVFLFPPKPKDTLMISRTPEVTCVVVDVSHEDPEVKFNWYVDGVEVHNAKTKPREEQYNSTYRVVSVLTVLHQDWLN<br>GKEYKCKVSNKALGAPIEKTISKAKGQPREPQVYTLPPSRDELTKNQVSLTCLVKGFYPSDIAVEWESNGQPENNYKTTPPVLDSDGSFFLYSKLTVDKS<br>RWQQGNVFSCSVMHEALHNHYTQKSLSLSPGK |
| Light chain* | RTVAAPSVFIFPPSDEQLKSGTASVVCLLNNFYPREAKVQWKVDNALQSGNSQESVTEQDSKDSTYSLSSTLTLSKADYEKHKVYACEVTHQGLSSPVT<br>KSFNRGEC |

*Exclude variable regions ($V_H$ or $V_L$)

**Figure S1.** Summary of antibody sequences. List of (A) antibody variable region sequences (VH and VL) and (B) sequences of heavy and light chains, excluding the variable regions.

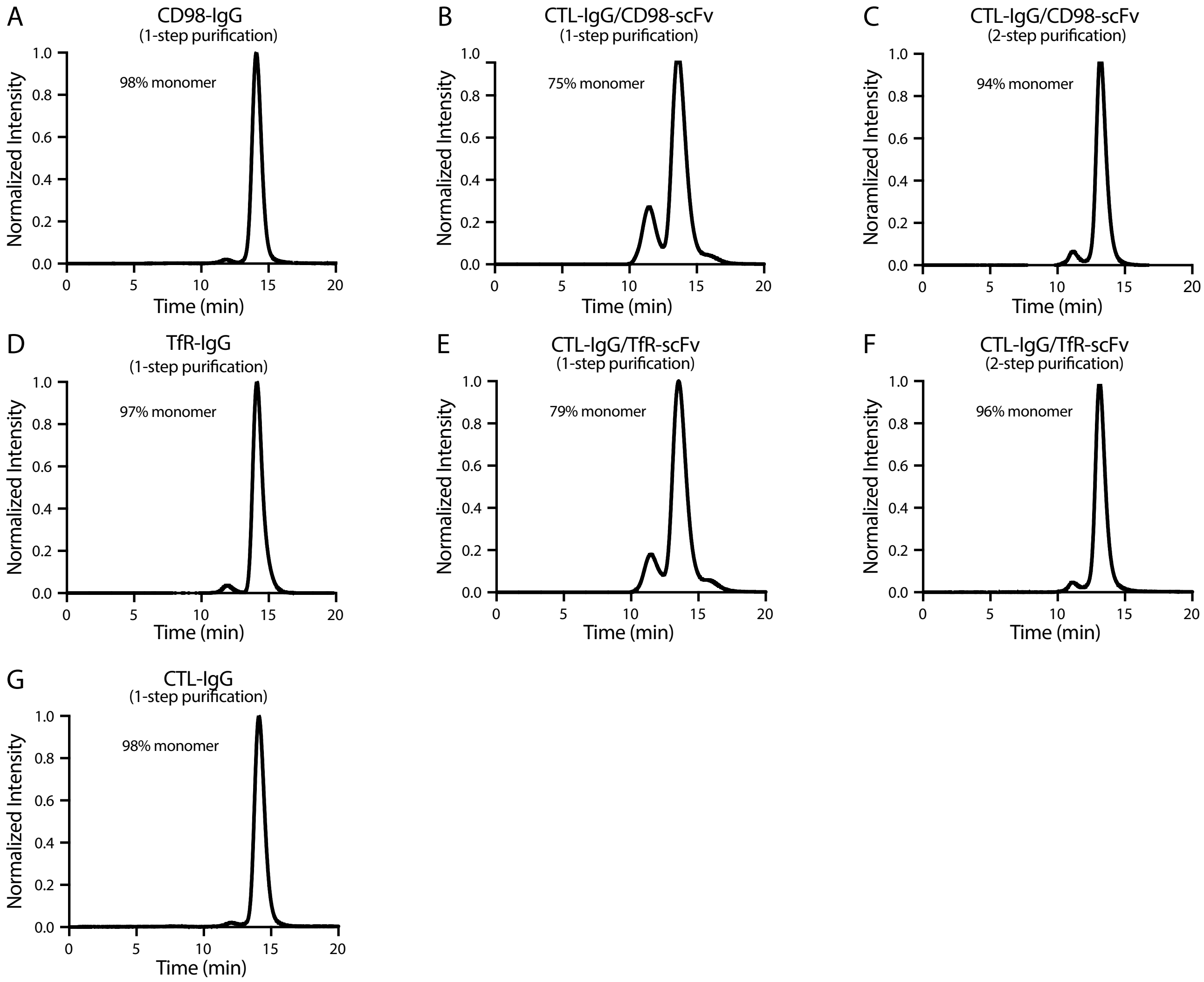


**Figure S2. Analytical size-exclusion chromatography analysis of antibodies produced in this work.** Representative chromatograms for (A) CD98-IgG (1-step purified), (B) CTL-IgG/CD98-scFv (1-step purified), (C) CTL-IgG/CD98-scFv (2-step purified), (D) TfR-IgG (1-step purified), (E) CTL-IgG/TfR-scFv (1-step purified), (F) CTL-IgG/TfR-scFv (2-step purified), and (G) CTL-IgG (1-step purified). All bispecific antibodies used in this work were 2-step purified to ensure >90% purity.

A

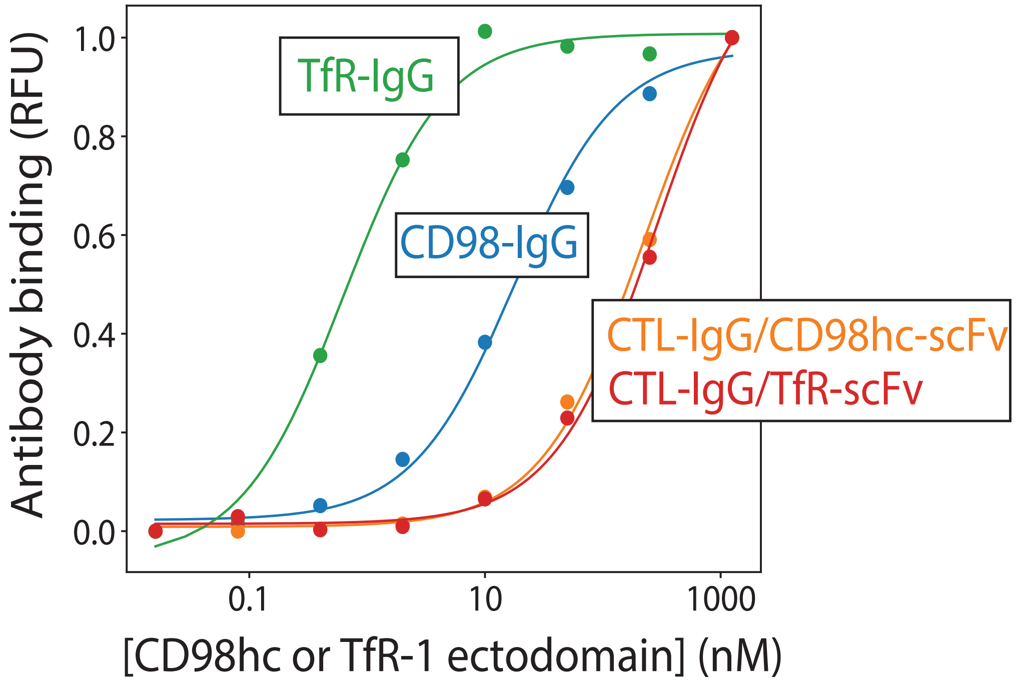


B

| | $EC_{50}$ ± Std. Dev. |
|---|---|
| TfR-IgG | 0.7±0.2 |
| CTL-IgG/TfR-scFv | 250±73 |
| CD98-IgG | 19.5±7.0 |
| CTL-IgG/CD98hc-scFv | 184±96 |

**Figure S3. Analysis of IgGs and bispecific antibodies binding to CD98hc and TfR-1ectodomains.** (A) Representative binding curves for the CD98hc and TfR-1 IgGs and bispecific antibodies. The antibodies were immobilized on Protein A Dynabeads, and their binding to biotinylated CD98hc and TfR-1 ectodomains was evaluated using flow cytometry. (B) Summary of binding measurements, which included three independent experiments. The values are averages, and the errors are standard deviations.

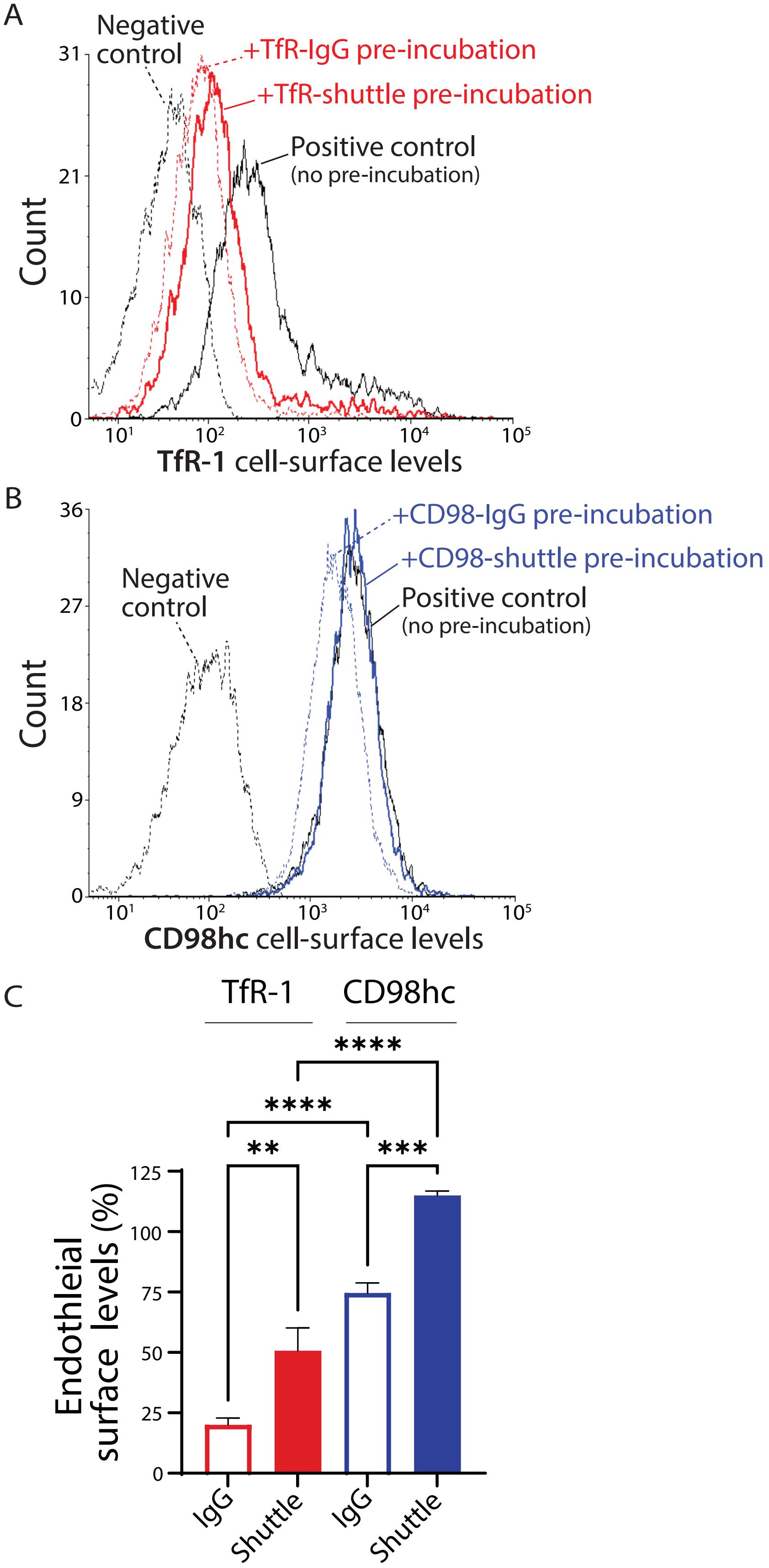


**Figure S4. Endothelial surface levels of CD98hc and TfR-1 after engagement with IgGs and bispecific antibody shuttles.** Mouse brain endothelial (bEnd.3) cells were incubated overnight with saturating concentrations of (A) TfR-1 and (B) CD98hc IgGs (100 nM) and shuttles (200 nM). Cell-surface levels were then assayed via flow cytometry using non-competitive antibodies. (C) Quantification of the endothelial surface levels of CD98hc and TfR-1 after incubation with IgGs and shuttles. In (A-C), the use of twice the shuttle concentration relative to the IgG concentration was to maintain a constant total number of antibody binding sites against CD98hc and TfR-1, as the shuttles are monovalent for binding CD98hc and TfR-1, while the IgGs are bivalent for binding these targets. The experiments were repeated multiple times (n=3), representative images are shown in (A) and (B), and normalized mean values and corresponding standard errors are shown in (C). p-values <0.01 (**), <0.001 (***), and <0.0001 (****) were determined by one-way ANOVA.

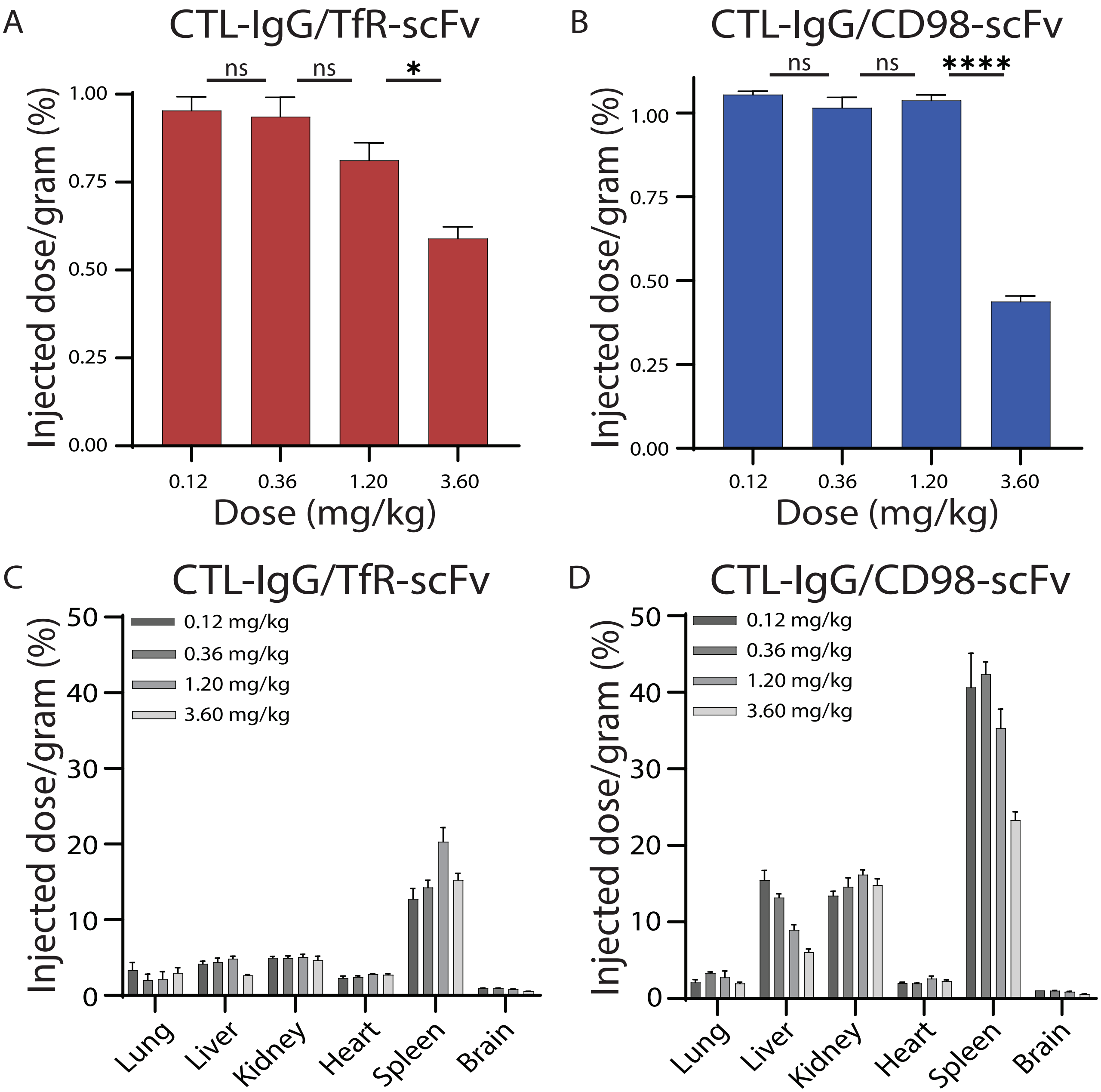


**Figure S5. Dose escalation analysis for the TfR-1 and CD98hc shuttles.** (A-D) The percent injected dose per gram (%ID/g) after 1 h in the brains at each dose for (A) CTL-IgG/TfR-scFv and (B) CTL-IgG/CD98-scFv, as well as in other organs for (C) CTL-IgG/TfR-scFv and (D) CTL-IgG/CD98-scFv. In (A-D), the shuttles were radiolabeled with 125I and injected retro-orbitally at a range of doses: 0.12, 0.36, 1.20, and 3.60 mg/kg (0.67, 2.0, 6.7, and 20.0 nmol/kg). Following transcardial perfusion with PBS at 1 h, the total organ %ID/g values were determined. n = 3-4 mice per group. p-values $<0.05$ (*) and $<0.0001$ (****) were determined by Welch's t-test.

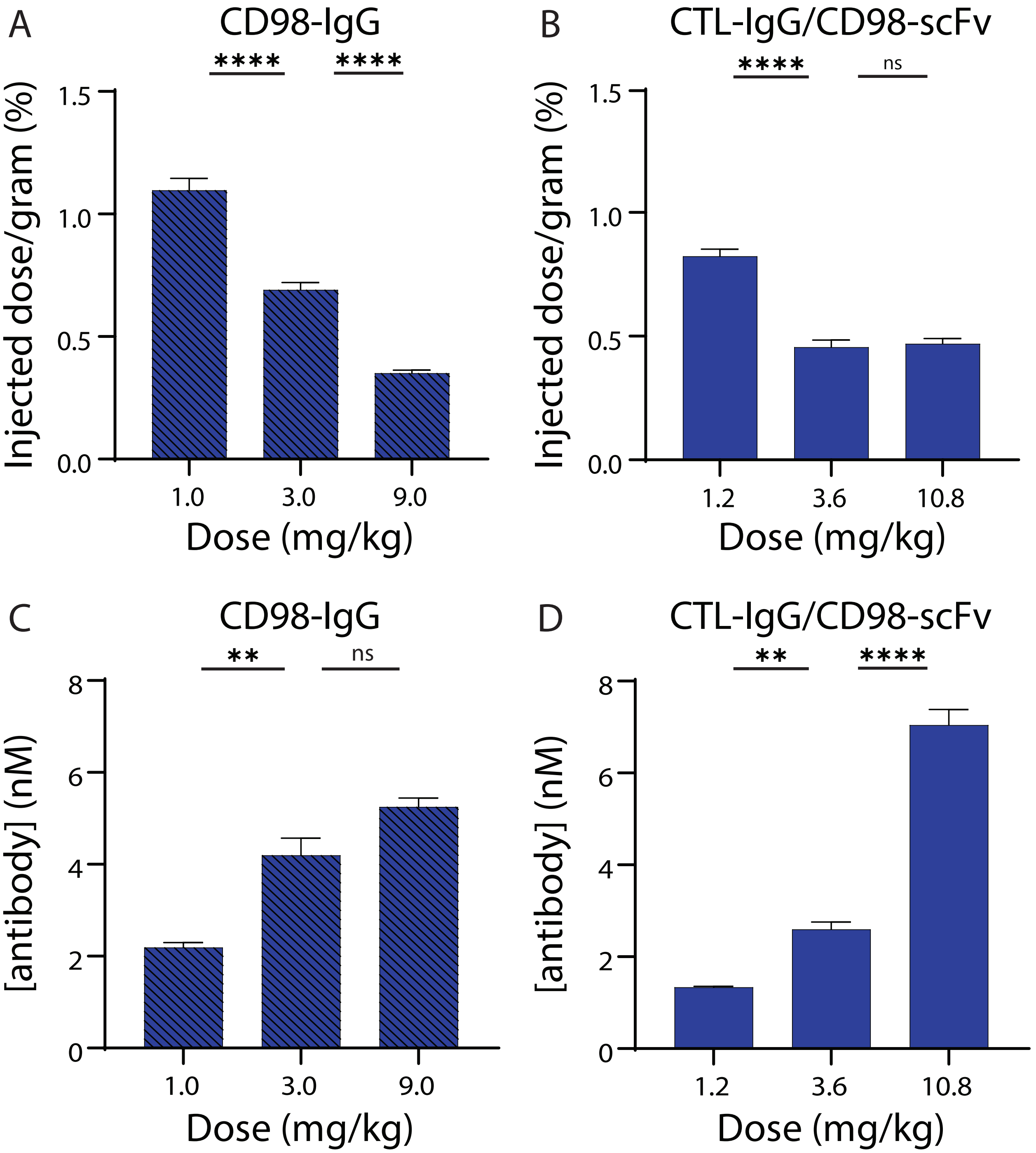


**Figure S6. Dose escalation analysis of CD98 mAb and shuttle.** CD98-IgG and and CTL-IgG/CD98-scFv were radiolabeled with $^{125}$I and injected retro-orbitally at a range of doses. Following transcardial perfusion with PBS at 24 h, percent injected dose per gram (%ID/g) and antibody concentration (nM) in the brains was evaluated. n = 4-5 mice per group. p-value <0.01 (**) and <0.0001 (****) determined by one-way ANOVA with multiple comparisons.

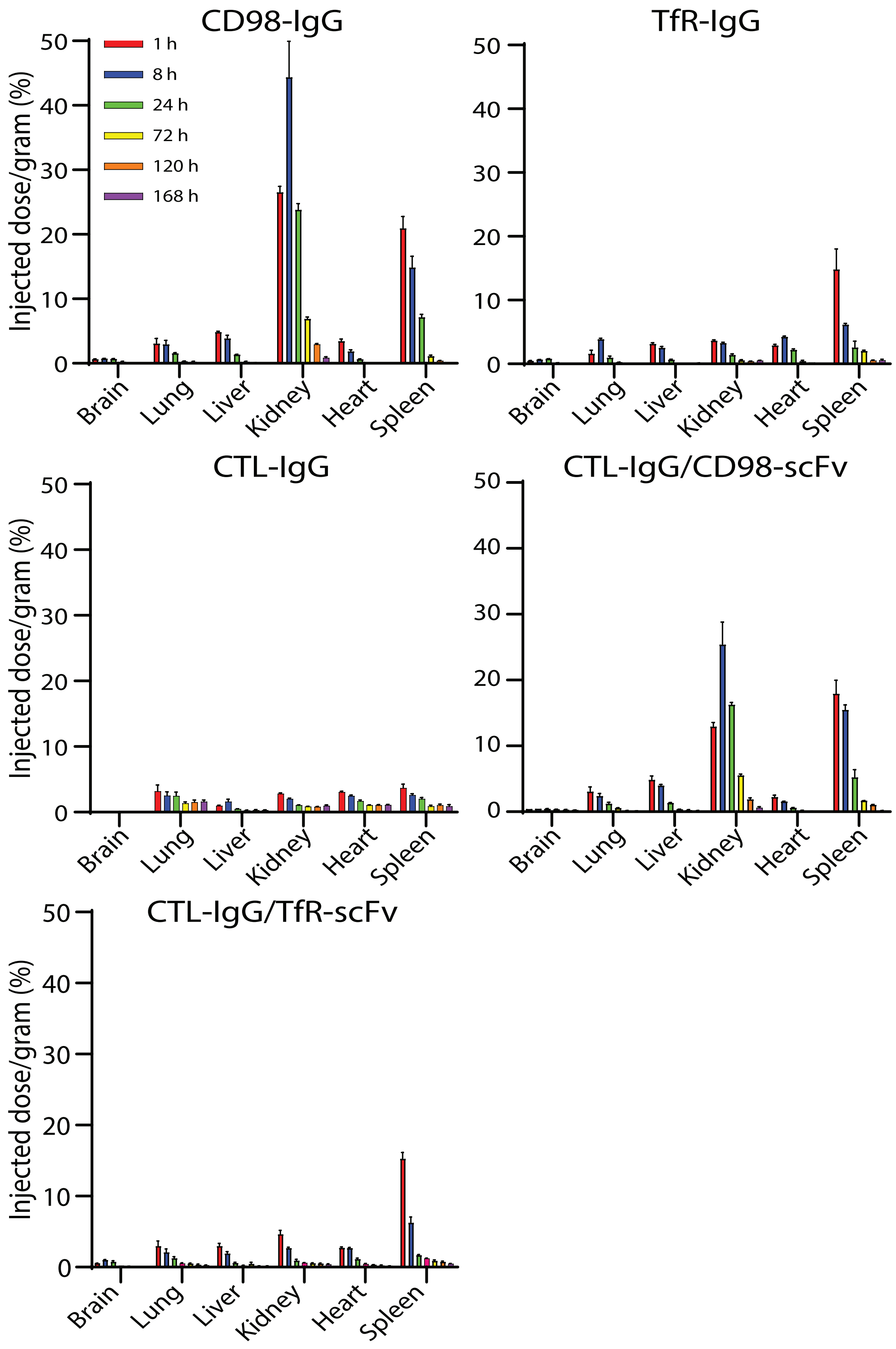


**Figure S7. Organ distribution of TfR-1 and CD98hc IgGs and shuttles.** Organ biodistribution, shown in %ID/g, of $^{125}I$ radiolabeled IgGs and shuttles (3 mg/kg for IgG and 3.6 mg/kg for shuttles; 20 nmol/kg) after retro-orbital injection for (A) CD98-IgG, (B) TfR-IgG, (C) CTL-IgG, (D) CTL-IgG/CD98-scFv, and (e) CTL-IgG/TfR-scFv. n=3-5 mice per group.

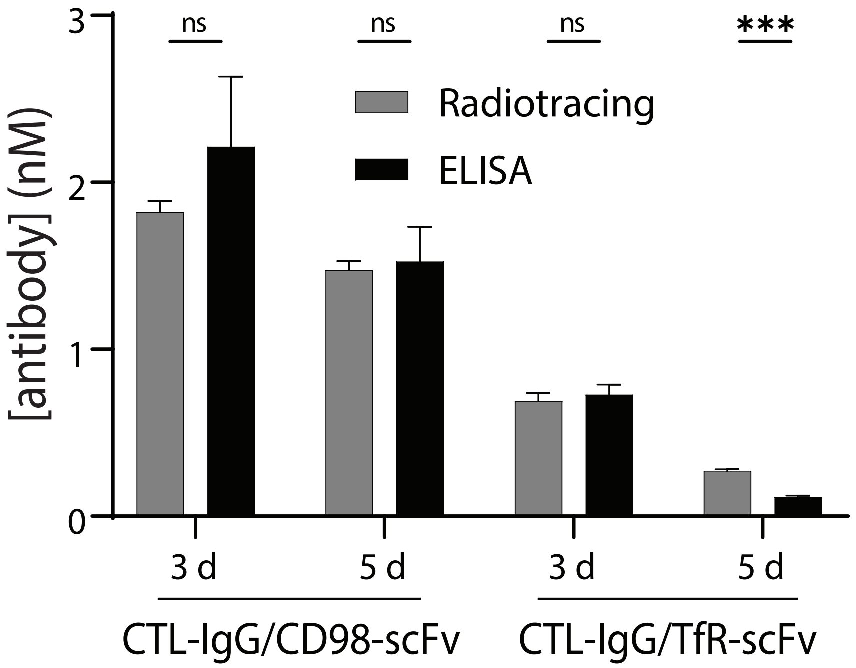


**Figure S8. Comparison of antibody brain concentrations using radiotracing and ELISA.** The brain concentrations of the shuttles (CTL-IgG/TfR-scFv and CTL-IgG/CD98-scFv) were analyzed by an IgG ELISA and compared to equivalent radiotracing ($^{125}I$) analysis. The injected dose was 3.6 mg/kg (20 nmol/kg). n=3-4 mice per group. p-values <0.001 (***) determined by unpaired t-tests.

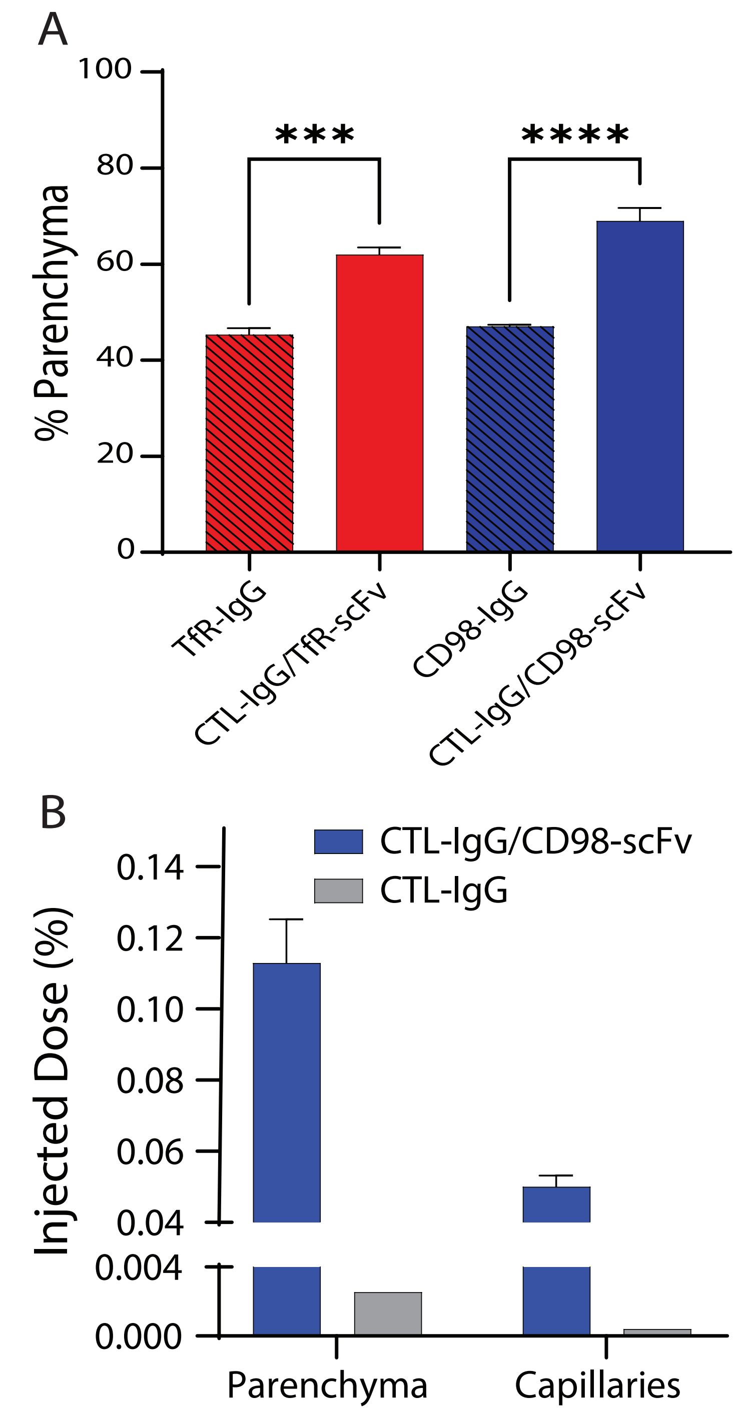


**Figure S9. Percentage of TfR-1 and CD98hc IgGs and shuttles in the brain parenchyma.** (A) Percentage of antibody in the brain parenchyma after 1 h. n=4 mice per group. p values <0.001 (***) and <0.0001 (****), as determined using a two-tailed t-test. (B) Percent of injected dose in the parenchyma and capillaries after 1 h. In (A) and (B), bispecific shuttles were given at a dose of 3.6 mg/kg and IgGs were given at a dose of 3 mg/kg (20 nmol/kg).

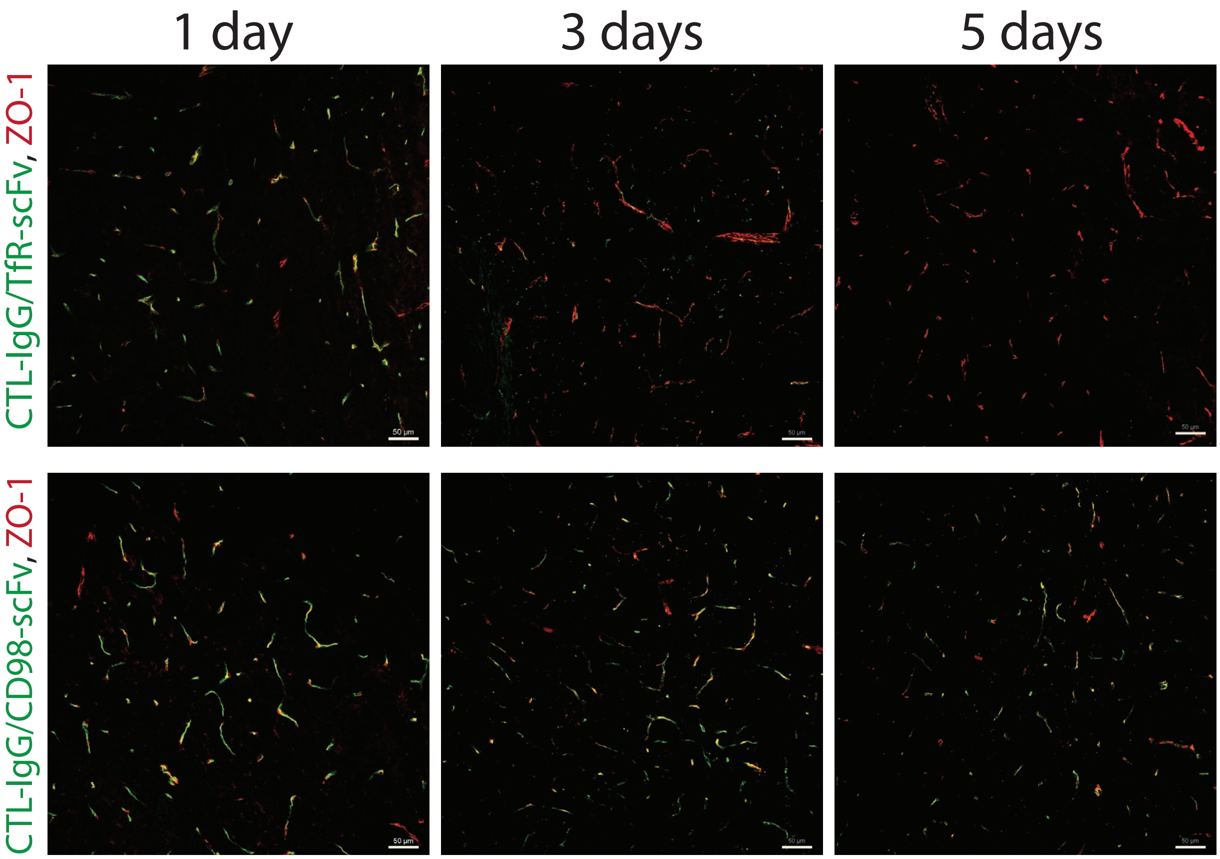


**Figure S10. Immunostaining of injected TfR-1 and CD98hc shuttles.** The shuttles (CTL-IgG/TfR-scFv and CTL-IgG/CD98-scFv) were labeled with Alexa Fluor-647 and administered to mice (3.6 mg/kg, 20 nmol/kg). Brains were harvested at 1, 3 and 5 days and sectioned for immunostaining. Injected antibody is shown in green and blood vessels (ZO-1) are shown in red. Scale bar is 50 µm.

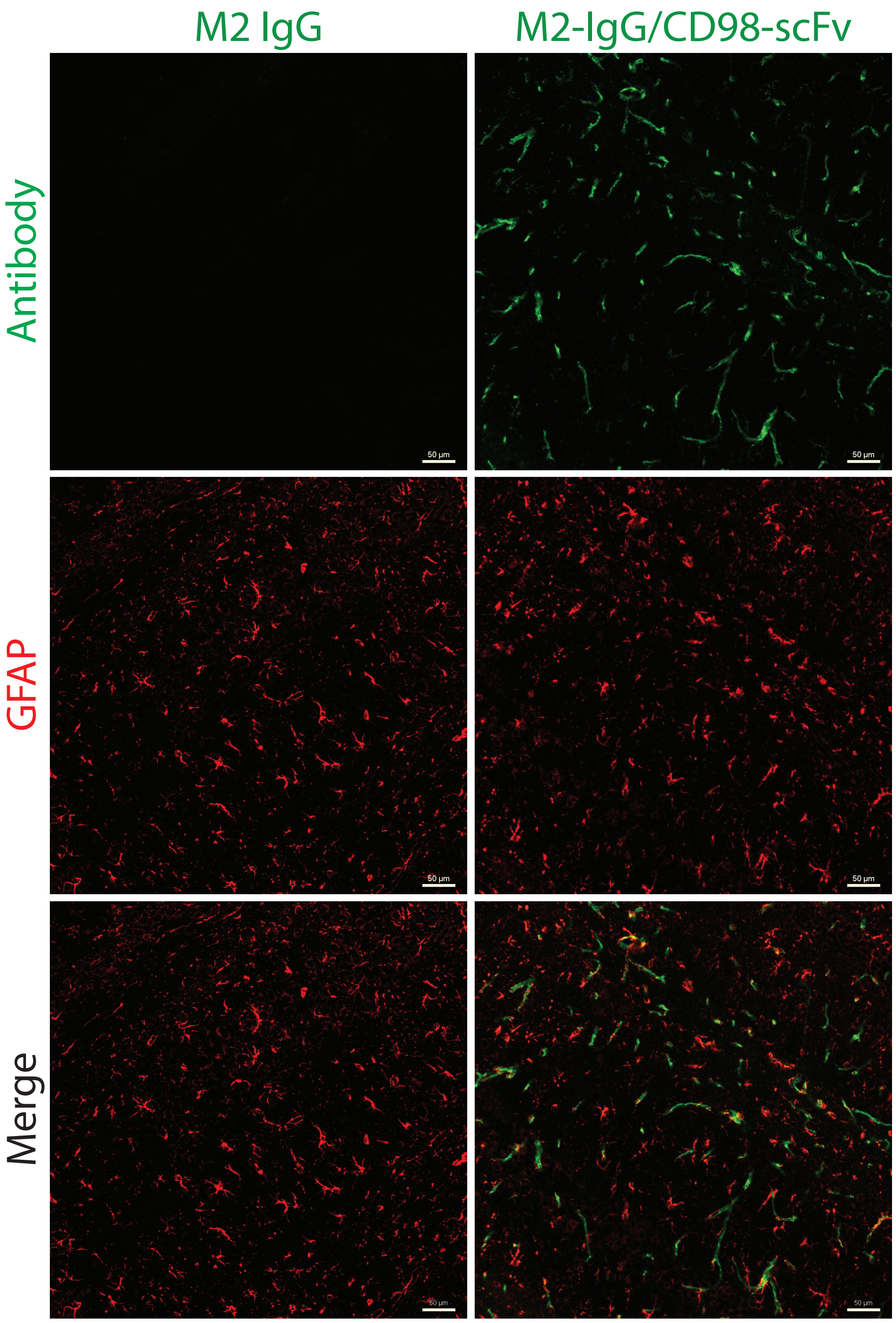


**Figure S11. Astrocyte-specific IgG is not detected in the brain without CD98hc-mediated shuttling.** Brain sections of mice injected with fluorescently labeled, astrocyte-specific IgG (M2) or M2-IgG/CD98-scFv corresponding to one day after intravenous administration. The injected antibodies (M2 and or M2-IgG/CD98-scFv) are shown in green relative to GFAP in red. Scale bar is 50 μm.

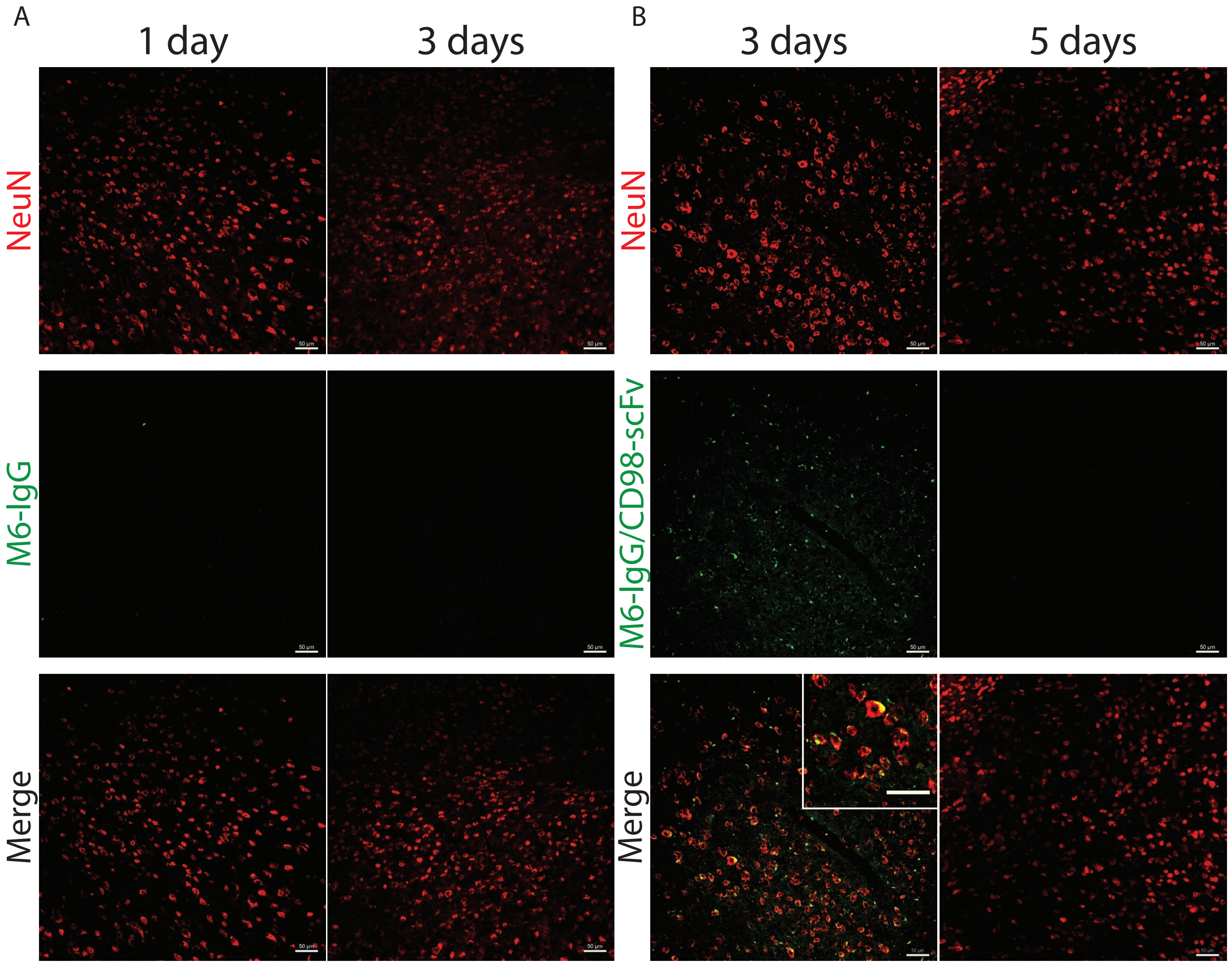


**Figure S12. Neuron-specific IgG is weakly detected in the brain without CD98hc-mediated shuttling.** Brain sections of mice injected with fluorescently labeled, neuron-specific (A) IgG (M6) or (B) M6-IgG/CD98-scFv corresponding to one day and/or three days after intravenous administration. The injected antibodies (M6 and or M6-IgG/CD98-scFv) are shown in green relative to NeuN+ neurons in red. Scale bar is 50 µm.

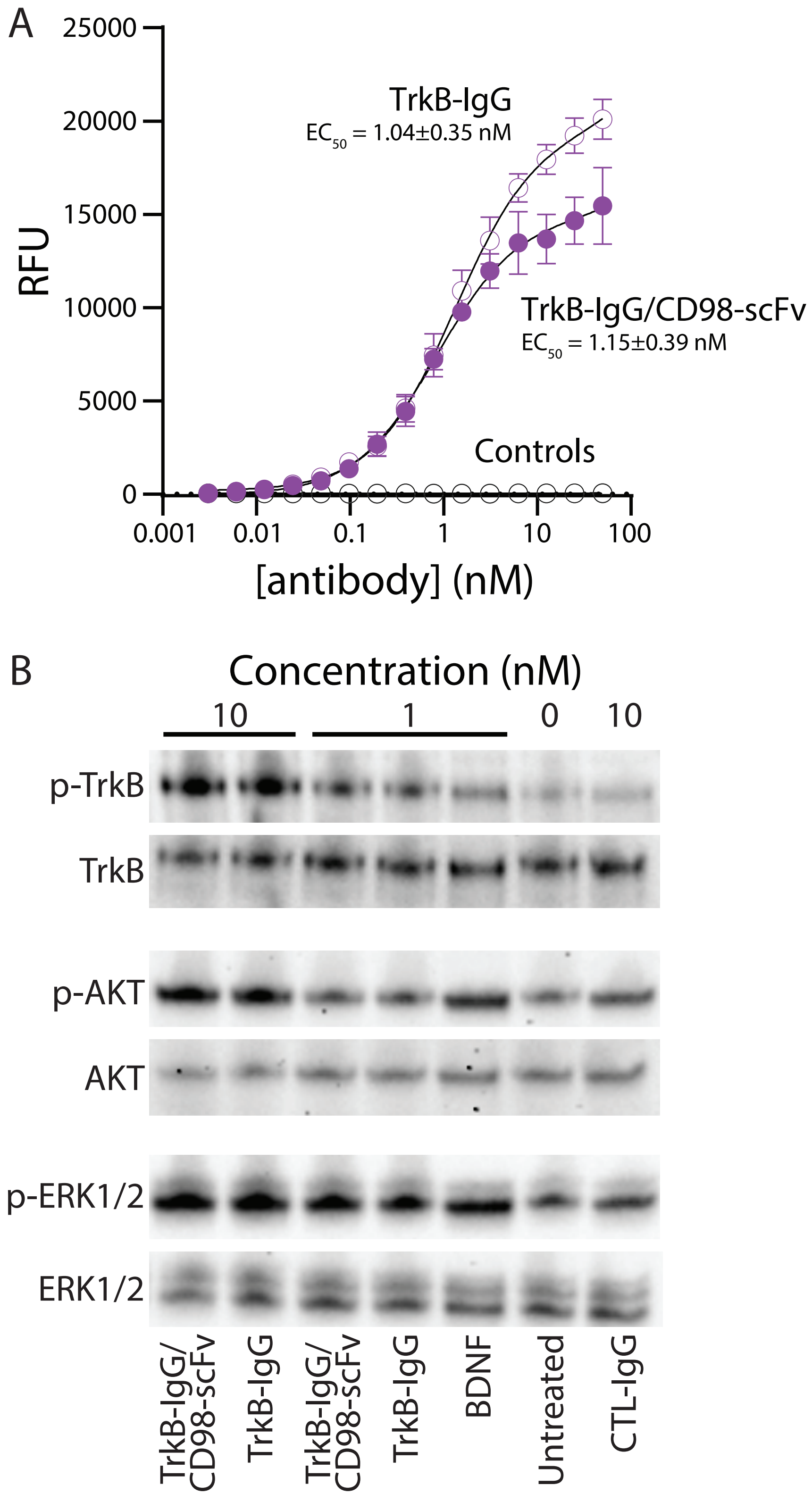


**Figure S13. In vitro characterization of TrkB IgG and TrkB/CD98hc shuttle.** (A) Concentration-dependent binding analysis of TrkB IgG and TrkB/CD98hc shuttle to REN cells expressing mouse TrkB. Controls were the same antibodies incubated with REN cells that do not express TrkB and a control IgG incubated with REN cells that express mouse TrkB. (B) Western blotting analysis of REN cells expressing mouse TrkB after incubation with antibodies (15 min). Cells were treated with BDNF (1 nM) as a positive control and a control IgG (10 nM) as a negative control.

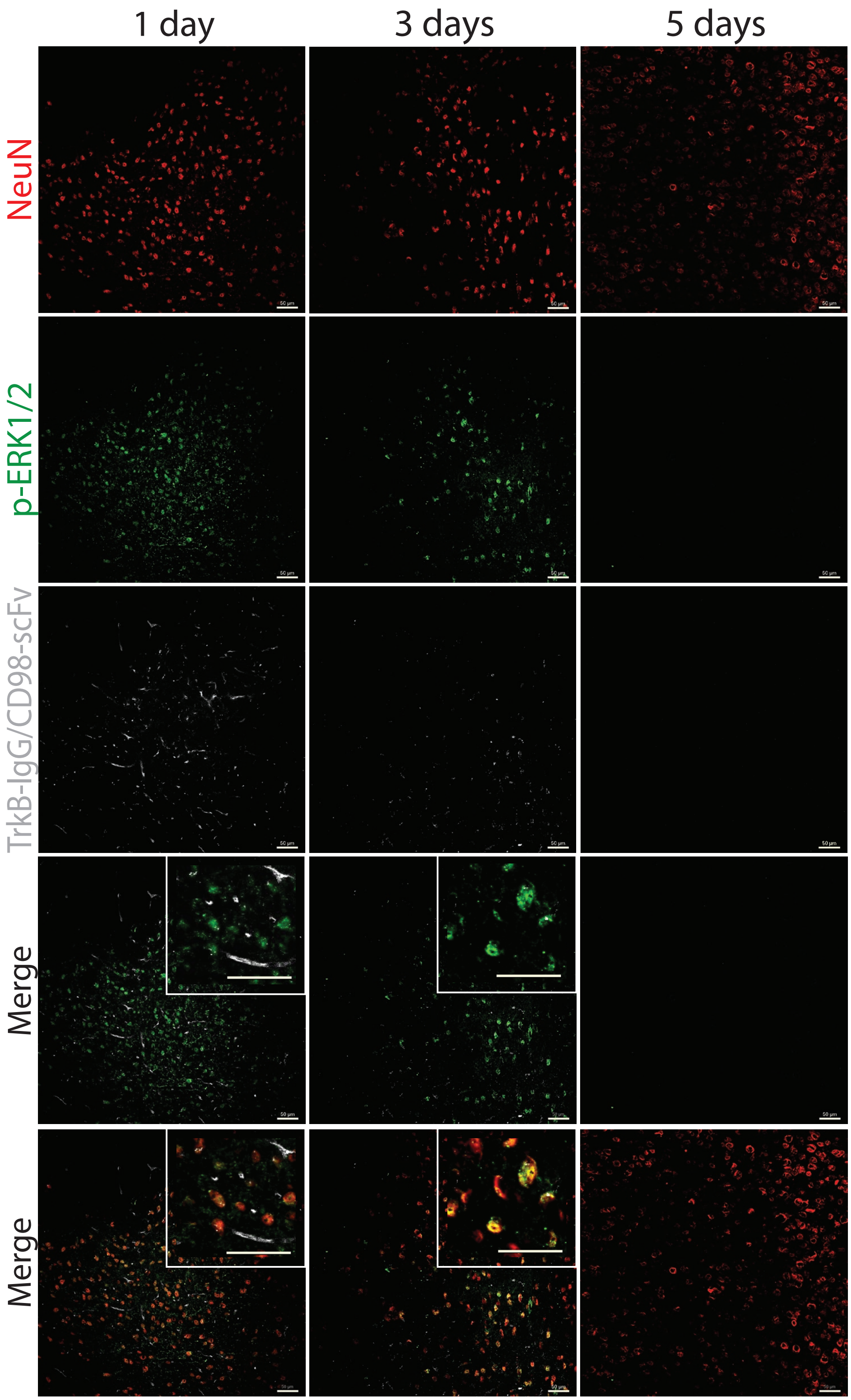


**Figure S14. Receptor activation (p-ERK1/2) is strongly detected for a CD98hc-shuttled TrkB agonist antibody.** Mouse brain sections 1-5 days after administration of TrkB/CD98hc shuttle. The injected antibody (TrkB/CD98hc shuttle, 3.6 mg/kg) is shown in grey relative to p-ERK1/2 in green and NeuN+ neurons in red. Scale bars are 50 µm.

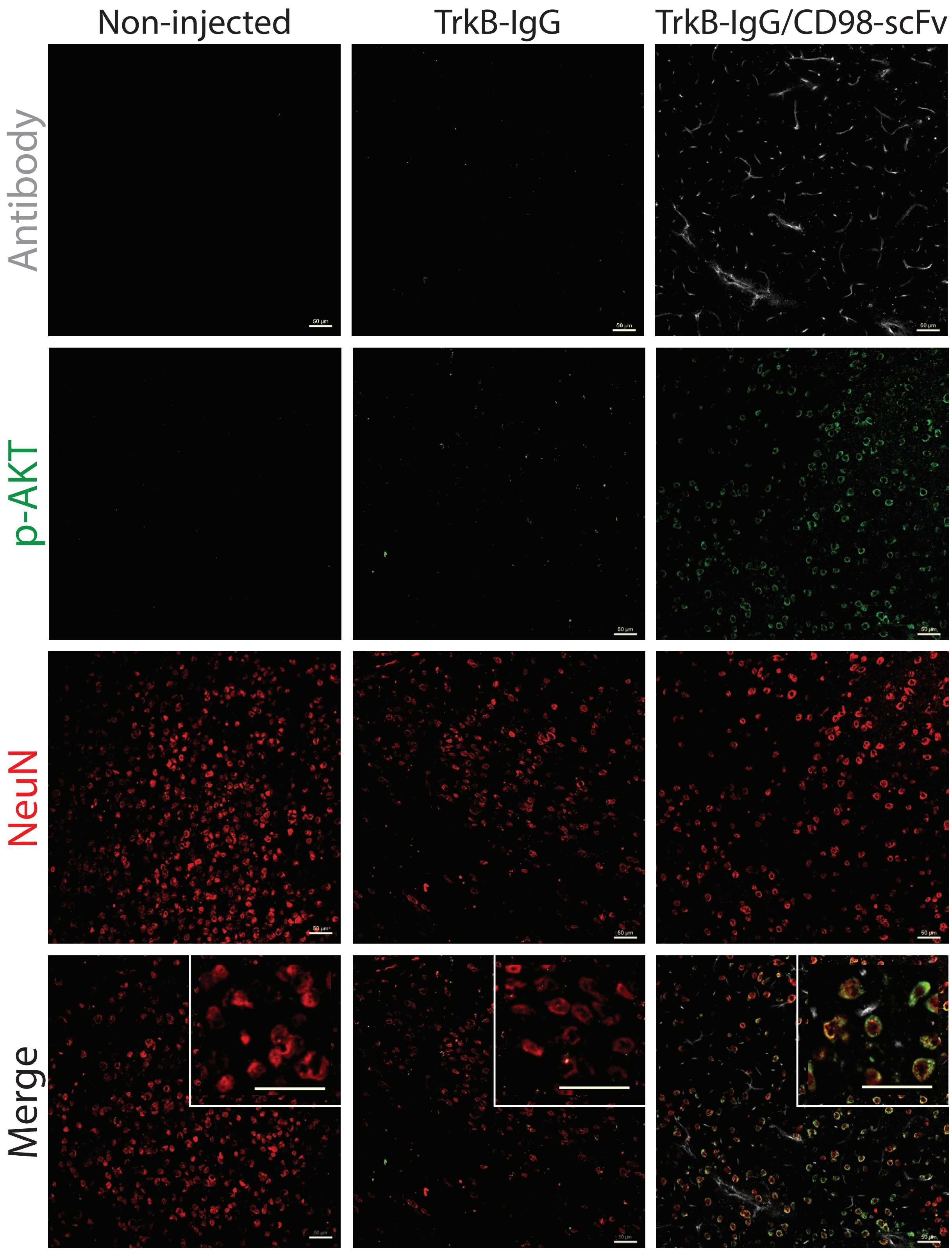


**Figure S15. Receptor activation (p-AKT) is weakly detected for a TrkB agonist antibody without CD98hc-mediated shuttling.** Mouse brain sections (left) without antibody administration (non-injected control), (center) one day after administration of TrkB, and (right) one day administration of TrkB/CD98hc shuttle. The injected antibodies (TrkB IgG and TrkB/CD98hc shuttle) are shown in grey relative to p-AKT in green and NeuN+ neurons in red. The antibody dose was 3 mg/kg for IgG and 3.6 mg/kg for shuttle (20 nmol/kg). Scale bars are 50 μm.

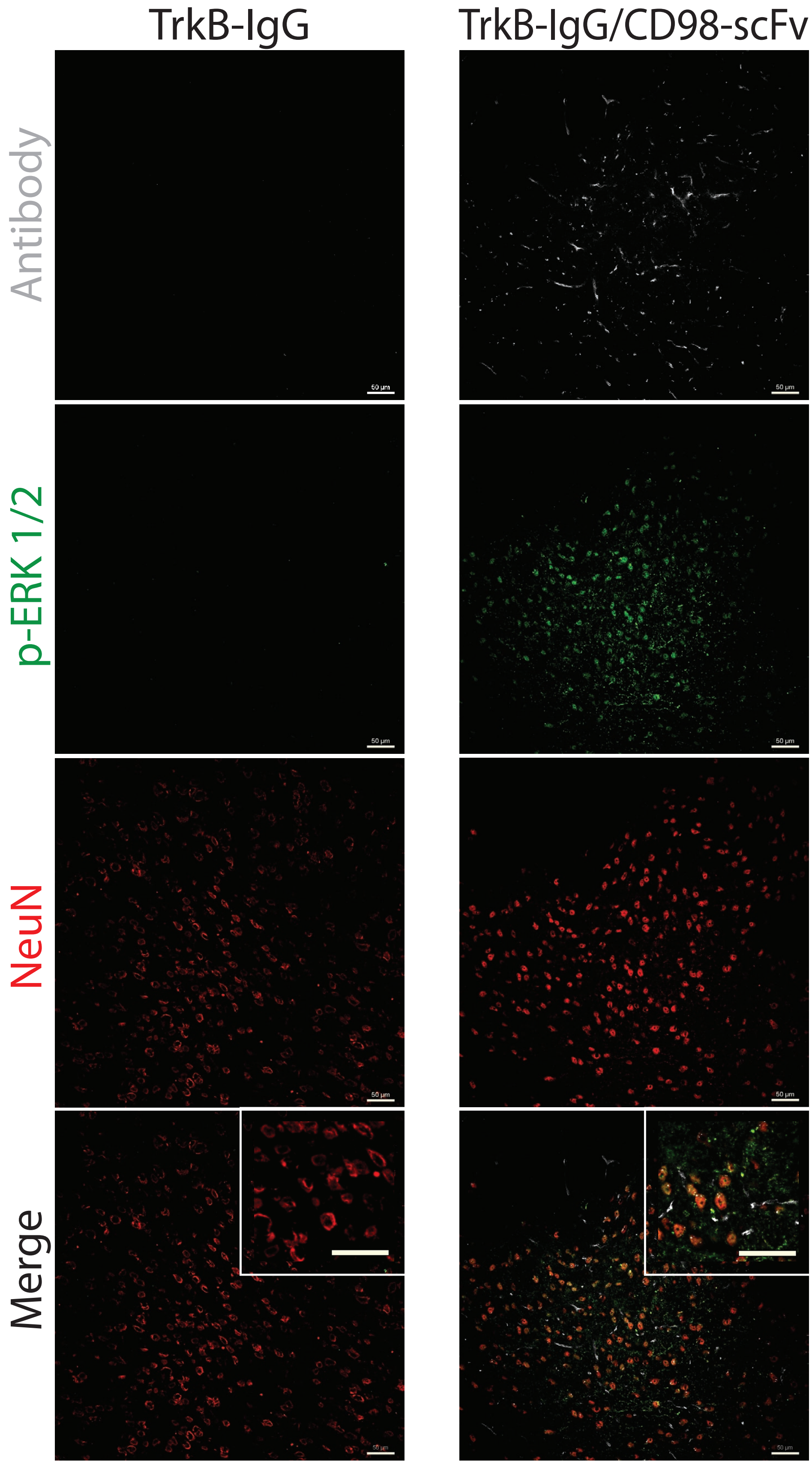


**Figure S16. Receptor activation (p-ERK1/2) is weakly detected for a TrkB agonist antibody without CD98hc-mediated shuttling.** Mouse brain sections one day after administration of (left) TrkB IgG and (right) TrkB/CD98hc shuttle. The injected antibodies are shown in grey relative to p-ERK1/2 in green and NeuN+ neurons in red. The antibody dose was 3 mg/kg for IgG and 3.6 mg/kg for shuttle (20 nmol/kg). Scale bars are 50 μm.

**Table S1.** Statistical comparison of brain antibody concentration by a one-way ANOVA at 1h, 8h, 24h, 72h, 120h, and 168h. p-Values were determined using Tukey significance test with $\alpha$ =.05.

| 1h | Summary | p-Value |
|---|---|---|
| CD98-IgG vs. CTL-IgG | **** | <0.0001 |
| CTL-IgG/CD98-scFv vs. CTL-IgG | **** | <0.0001 |
| CTL-IgG/TfR-scFv vs. CTL-IgG | **** | <0.0001 |
| TfR-IgG vs. CTL-IgG | **** | <0.0001 |
| CD98-IgG vs. CTL-IgG/CD98-scFv | *** | 0.0002 |
| CTL-IgG/TfR-scFv vs. TfR-IgG | * | 0.0486 |
| CD98-IgG vs. TfR-IgG | ns | 0.6383 |
| CTL-IgG/CD98-scFv vs. CTL-IgG/TfR-scFv | **** | <0.0001 |
| **8h** | | |
| CD98-IgG vs. CTL-IgG | **** | <0.0001 |
| CTL-IgG/CD98-scFv vs. CTL-IgG | **** | <0.0001 |
| CTL-IgG/TfR-scFv vs. CTL-IgG | **** | <0.0001 |
| TfR-IgG vs. CTL-IgG | **** | <0.0001 |
| CD98-IgG vs. CTL-IgG/CD98-scFv | **** | <0.0001 |
| CTL-IgG/TfR-scFv vs. TfR-IgG | **** | <0.0001 |
| CD98-IgG vs. TfR-IgG | ns | 0.1644 |
| CTL-IgG/CD98-scFv vs. CTL-IgG/TfR-scFv | **** | <0.0001 |
| **24h** | | |
| CD98-IgG vs. CTL-IgG | **** | <0.0001 |
| CTL-IgG/CD98-scFv vs. CTL-IgG | *** | 0.0006 |
| CTL-IgG/TfR-scFv vs. CTL-IgG | **** | <0.0001 |
| TfR-IgG vs. CTL-IgG | **** | <0.0001 |
| CD98-IgG vs. CTL-IgG/CD98-scFv | *** | 0.0003 |
| CTL-IgG/TfR-scFv vs. TfR-IgG | ns | >0.9999 |
| CD98-IgG vs. TfR-IgG | ns | 0.9954 |
| CTL-IgG/CD98-scFv vs. CTL-IgG/TfR-scFv | *** | 0.0009 |
| **72h** | | |
| CD98-IgG vs. CTL-IgG | **** | <0.0001 |
| CTL-IgG/CD98-scFv vs. CTL-IgG | **** | <0.0001 |
| CTL-IgG/TfR-scFv vs. CTL-IgG | ** | 0.0024 |
| TfR-IgG vs. CTL-IgG | **** | <0.0001 |
| CD98-IgG vs. CTL-IgG/CD98-scFv | ns | 0.7962 |
| CTL-IgG/TfR-scFv vs. TfR-IgG | * | 0.0128 |
| CD98-IgG vs. TfR-IgG | **** | <0.0001 |
| CTL-IgG/CD98-scFv vs. CTL-IgG/TfR-scFv | **** | <0.0001 |
| **120h** | | |
| CD98-IgG vs. CTL-IgG | **** | <0.0001 |

| | | |
|---|---|---|
| CTL-IgG/CD98-scFv vs. CTL-IgG | **** | <0.0001 |
| CTL-IgG/TfR-scFv vs. CTL-IgG | ns | 0.8691 |
| TfR-IgG vs. CTL-IgG | ns | 0.9997 |
| CD98-IgG vs. CTL-IgG/CD98-scFv | **** | <0.0001 |
| CTL-IgG/TfR-scFv vs. TfR-IgG | ns | 0.7367 |
| CD98-IgG vs. TfR-IgG | **** | <0.0001 |
| CTL-IgG/CD98-scFv vs. CTL-IgG/TfR-scFv | **** | <0.0001 |
| 168h | | |
| CD98-IgG vs. CTL-IgG | **** | <0.0001 |
| CTL-IgG/CD98-scFv vs. CTL-IgG | **** | <0.0001 |
| CTL-IgG/TfR-scFv vs. CTL-IgG | *** | 0.0007 |
| TfR-IgG vs. CTL-IgG | *** | 0.0005 |
| CD98-IgG vs. CTL-IgG/CD98-scFv | **** | <0.0001 |
| CTL-IgG/TfR-scFv vs. TfR-IgG | ns | 0.9803 |
| CD98-IgG vs. TfR-IgG | **** | <0.0001 |
| CTL-IgG/CD98-scFv vs. CTL-IgG/TfR-scFv | **** | <0.0001 |

**Table S2. Summary of antibodies used in this work for brain immunostaining and western blotting.**

| Antibody | Dilution | Application | Vendor | Catalog Number |
|---|---|---|---|---|
| NeuN | 1:200 | Immunostaining | Thermo Fisher | MA5-33103 |
| GFAP | 1:200 | Immunostaining | Thermo Fisher | PA1-10019 |
| pAKT | 1:100 | Immunostaining | Thermo Fisher | 44-621G |
| pERK | 1:200 | Immunostaining | Thermo Fisher | 36-8800 |
| ZO-1 | 1:100 | Immunostaining | Thermo Fisher | 339188 |
| AKT | 1:2000 | Western blotting | Cell Signaling Technology (CST) | 4685 |
| ERK | 1:2000 | Western blotting | CST | 9102 |
| TrkB | 1:1000 | Western blotting | CST | 4603 |
| pAKT | 1:1000 | Western blotting | CST | 4060 |
| pERK | 1:1000 | Western blotting | CST | 9101 |
| pTrkB | 1:1000 | Western blotting | Thermo Fisher | PA5-36695 |
| Anti-rabbit mAb HRP conjugated | 1:3000 | Western blotting | CST | 7074 |
